# Adaptation, ancestral variation and gene flow in a ‘Sky Island’ *Drosophila* species

**DOI:** 10.1101/2020.05.14.096008

**Authors:** Tom Hill, Robert L. Unckless

## Abstract

Over time, populations of species can expand, contract, fragment and become isolated, creating subpopulations that must adapt to local conditions. Understanding how species maintain variation after divergence as well as adapt to these changes in the face of gene flow, is of great interest, especially as the current climate crisis has caused range shifts and frequent migrations for many species. Here, we characterize how a mycophageous fly species, *Drosophila innubila*, came to inhabit and adapt to its current range which includes mountain forests in southwestern USA separated by large expanses of desert. Using population genomic data from more than 300 wild-caught individuals, we examine four populations to determine their population history in these mountain forests, looking for signatures of local adaptation. We find *D. innubila* spread northwards during the previous glaciation period (30-100 KYA), and has recently expanded even further (0.2-2 KYA). *D. innubila* shows little evidence of population structure, consistent with a recent establishment and genetic variation maintained since before geographic stratification. We also find some signatures of recent selective sweeps in chorion proteins and population differentiation in antifungal immune genes suggesting differences in the environments to which flies are adapting. However, we find little support for long-term recurrent selection in these genes. In contrast, we find evidence of long-term recurrent positive selection in immune pathways such as the Toll-signaling system and the Toll-regulated antimicrobial peptides.

## Introduction

In the past 25,000 years, the earth has undergone substantial environmental changes due to both human-mediated events (anthropogenic environment destruction, desert expansion, extreme weather and the growing anthropogenic climate crisis) (Cloudsley-Thompson 1978; Rosenzweig *et al*. 2008) and events unrelated to humans (glaciation and tectonic shifts) (Hewitt 2000; Holmgren *et al*. 2003; Arizona-Geological-Survey 2005). These environmental shifts can fundamentally reorganize habitats, and influence organism fitness, rates of migration between locations, and population ranges (Smith *et al*. 1995; Astanei *et al*. 2005; Rosenzweig *et al*. 2008; Searle *et al*. 2009; Cini *et al*. 2012; Porretta *et al*. 2012; Antunes *et al*. 2015). Signatures of the how organisms adapt to these events, or to population migration, are often etched in the patterns of molecular variation within and between species (Charlesworth *et al*. 2003; Wright *et al*. 2003; Excoffier *et al*. 2009).

The Madrean archipelago is an environment currently undergoing extensive environmental change due to the climate crisis, and presents a good model environment to study the genomic consequences of local adaptation and gene flow in a changing environment (Smith and Farrell 2005; Coe *et al*. 2012; Manthey and Moyle 2015; Wiens *et al*. 2019). This range, located in southwestern USA and northwestern Mexico, contains numerous forested mountains known as ‘Sky islands’, separated by large expanses of desert (McCormack *et al*. 2009; Coe *et al*. 2012). These ‘islands’ were connected by woodland forests during the previous glacial maximum which then retreated, leaving montane forest habitat separated by hundreds of miles of desert, presumably limiting migration between locations for most species (Arizona-Geological-Survey 2005; Smith and Farrell 2005; McCormack *et al*. 2009). The islands are hotbeds of ecological diversity. However, due to the changing climate in the past 100 years, they have become more arid and prone to wild fires, which may drive migration, adaptation and even extinction events (McCormack *et al*. 2009; Coe *et al*. 2012; Fave *et al*. 2015; Manthey and Moyle 2015).

Several studies of genetic diversity in Sky Island populations have explored the how species persist in these environments and the extent that gene flow occurs between locations. These studies primarily find that while birds are mostly unstructured (McCormack *et al*. 2009; Manthey and Moyle 2015), flightless species are highly structured (Smith and Farrell 2005; McCormack *et al*. 2009; Fave *et al*. 2015; Wiens *et al*. 2019), due to the difficulty in migrating between these mountains separated by hundreds of miles of desert. These studies also highlight that many species are adapting to the increasingly hot and arid conditions of the Sky Islands, caused by climate change (McCormack *et al*. 2009; Coe *et al*. 2012; Misztal *et al*. 2013). Specifically, several animal species are changing their population distributions in the Sky Islands and adapting to increasingly hostile conditions (Fave *et al*. 2015; Wiens *et al*. 2019). Additionally, the delay in the monsoon period has resulted in several plant species pushing back their flowering period, until adequate water is available (McCormack *et al*. 2009; Crimmins *et al*. 2011).

To date, no studies have considered demographics and local adaptation in the Sky Islands from a whole genome perspective and few have considered demography and local adaptation in insects (Fave *et al*. 2015). Luckily, the Madrean archipelago is inhabited by the ecological *Drosophila* model species, *Drosophila innubila* (Jaenike et al. 2003; Dyer and Jaenike 2005). *Drosophila innubila* is a mycophageous species found throughout these Sky islands and thought to have arrived during the last glacial maximum (Dyer and Jaenike 2005; Dyer *et al*. 2005). This ‘island’ endemic mushroom-feeder emerges during the rainy season in the Sky Islands for 10-12 weeks in late summer and early fall, before returning to diapause for the winter and dry seasons (Patterson 1954). *D. innubila* and other mycophageous *Drosophila* are well-studied ecologically, giving us a thorough understanding of their life history, environment and pathogens (Shoemaker *et al*. 1999; Perlman *et al*. 2003; Dyer and Jaenike 2005; Dyer *et al*. 2005; Jaenike and Dyer 2008; Unckless 2011; Unckless and Jaenike 2011). For example, *D. innubila* is infected with a male-killing, maternally transmitted pathogen, called *Wolbachia* which has been extensively studied (Jaenike *et al*. 2003; Dyer 2004; Dyer and Jaenike 2005; Dyer *et al*. 2005). *D. innubila* is also frequently exposed to a highly virulent DNA virus (Unckless 2011), as well as toxic mushrooms (Scott Chialvo and Werner 2018). Given these environmental and pathogenic factors, we expect to identify recent signatures of evolution in the *D. innubila* genome on establishment in the Sky Islands, as well as signatures of a recent population migration.

Newly established populations are almost always small because they usually consist of a few founders (Charlesworth *et al*. 2003; Excoffier *et al*. 2009; Li and Durbin 2011). This results in a loss of rare alleles in the population (Tajima 1989; Gillespie 2004). The population will then grow to fill the carrying capacity of the new niche and adapt to the unique challenges in the new environment, both signaled by an excess of rare alleles (Excoffier *et al*. 2009; White *et al*. 2013). This adaptation can involve selective sweeps from new mutations or standing genetic variation, and signatures of adaptive evolution and local adaptation in genes key to the success of the population in this new location (Charlesworth *et al*. 2003; Hermisson and Pennings 2005; McVean 2007; Messer and Petrov 2013). However, these signals can confound each other making inference of population history difficult. For example, both population expansions and adaptation lead to an excess of rare alleles, meaning more thorough analysis is required to identify the true cause of the signal (Wright *et al*. 2003). Additionally, continual migration between populations can alter the allele frequencies within populations, possibly removing adaptive alleles, or supplying beneficial alleles from other subpopulations (Tigano and Friesen 2016). Signatures of demographic change are frequently detected in species that have recently undergone range expansion due to human introduction (Astanei *et al*. 2005; Excoffier *et al*. 2009) or the changing climate (Hewitt 2000; Parmesan and Yohe 2003; Guindon *et al*. 2010; Walsh *et al*. 2011; Cini *et al*. 2012). Other hallmarks of a range expansion include signatures of bottlenecks visible in the site frequency spectrum, and differentiation between populations (Charlesworth *et al*. 2003; Li and Durbin 2011). This can be detected by a deficit of rare variants, a decrease in population pairwise diversity and an increase in the statistic, Tajima’s D (Tajima 1989). Following the establishment and expansion of a population, there is an excess of rare variants and local adaptation results in divergence between the invading population and the original population. These signatures are also frequently utilized in human populations to identify traits which have fixed upon the establishment of human populations in a new location, or to identify how our human ancestors spread globally (Li and Durbin 2011).

Populations with limited migration provide a rare opportunity to observe replicate bouts of evolutionary change and this is particularly interesting regarding coevolution with pathogens, as could be examined in *D. innubila* (Dyer and Jaenike 2005; Unckless 2011).We sought to reconstruct the demographic and migratory history of *D. innubila* inhabiting the Sky islands to understand if migration is occurring between Sky Islands, and how the species is adapting to its new environment, based on signatures of selection in the genome (Tigano and Friesen 2016). Additionally, we hoped to determine if populations are locally adapting to their recent environment or if signatures of local adaptation are swamped out by species-level adaptation occurring before population divergence. We resequenced whole genomes of wild-caught individuals from four populations of *D. innubila* in four different Sky island mountain ranges. This differs from most population genomic studies of *Drosophila*, which rely on lab maintained inbred strains. Interestingly, we find little evidence of population structure by location, with structure limited to the mitochondria and a single autosome. However, we do find some signatures of local adaptation, such as for cuticle development and fungal pathogen resistance. We also find evidence of mitochondrial translocations into the nuclear genome, with strong evidence of local adaptation of these translocations, suggesting potential adaptation to changes in metabolic process of the host between location (Jaenike and Dyer 2008). Finally, we find signatures of long-term selection of the Toll immune pathway, across all populations.

## Results

### Drosophila innubila has recently expanded its geographic range and shows little divergence between geographically isolated populations

To characterize how *D. innubila* came to inhabit its current range, we collected flies from four Sky island locations across Arizona in September 2017: Chiricahuas (CH, 81 flies), Huachucas (HU, 48 flies), Prescott (PR, 84 flies) and Santa Ritas (SR, 67 flies) (Locations shown in Figure 1). Interestingly, while previous surveys mostly failed to collect *D. innubila* north of the Madrean archipelago in Prescott (Dyer and Jaenike 2005), we easily sampled from that location, suggesting a possible recent colonization (though we were also unable to collect *D. innubila* in the exact locations previously sampled) (Dyer and Jaenike 2005). If this was a recent colonization event, it could be associated with the changing climate of the area leading to conditions more accommodating to *D. innubila*.

**Figure 1:**
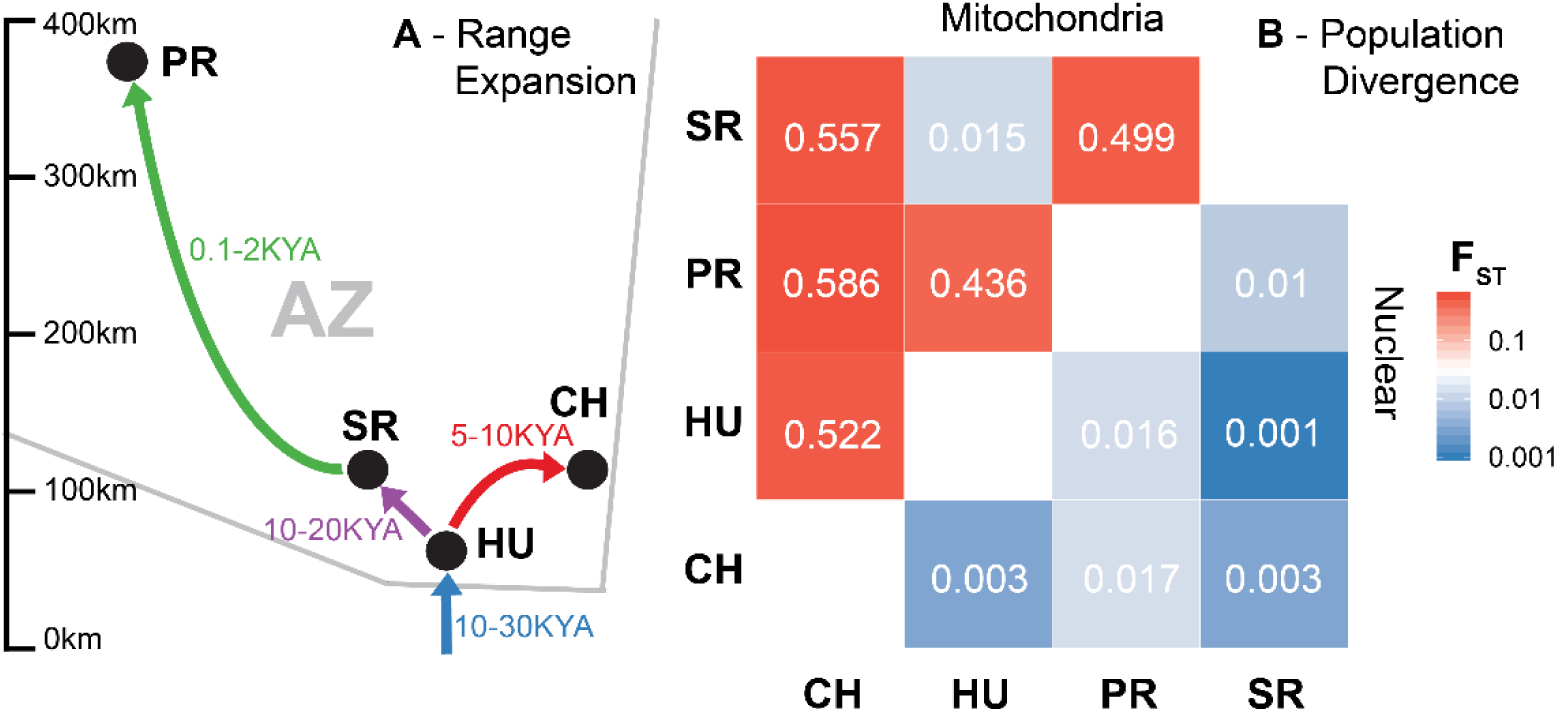
**A**. Schematic of the route of colonization of *D. innubila*, inferred from population size history based on StairwayPlot results across the four sample locations in Arizona (AZ), Chiracahua’s (CH, red), Huachucas (HU, blue), Prescott (PR, green) and Santa Ritas (SR, purple). Stairway plot results used to infer this history are shown in Supplementary Figure 1. **B**. Median pairwise F_ST_ between populations, comparing the autosomal nuclear genome (below diagonal) and the mitochondrial genome (above diagonal).

To determine when *D. innubila* established in each location and rates of migration between locations, we isolated and sequenced the DNA from our sampled *D. innubila* populations and characterized genomic variation. We then examined the population structure and changes in demographic history of *D. innubila* using silent polymorphism in StairwayPlot (Liu and Fu 2015). We found all sampled populations have a current estimated effective population size (N_e_) of ∼1 million individuals and an ancestral N_e_ of ∼4 million individuals, and all experience a bottleneck between 70 and 100 thousand years ago to an N_e_ of 10-20 thousand (Figure 1, Supplementary Figure 1A & B). This bottleneck coincides with a known glaciation period occurring in Arizona (Arizona-Geological-Survey 2005). Each surveyed population then appears to go through a population expansion between one and thirty thousand years ago, with populations settling from south to north (Figure 1A, Supplementary Figure 1A & B). Specifically, while the Huachucas population appears to have been colonized first (10-30 thousand years ago), the Prescott population was colonized much more recently (200-2000 years ago). This, and the absence of *D. innubila* in Prescott sampling until ∼2016 sampling suggests very recent northern expansion of *D. innubila* (Figure 1). Note, however, that StairwayPlot (Liu and Fu 2015) has estimated large error windows for Prescott, meaning the colonization could be more recent or ancient than the 200-2000 year estimate.

The *Drosophila* genome is organized into Muller elements, these are 6 chromosome arms which are generally conserved across all *Drosophila* species, named A to F (with A representing the ancestral *Drosophila* X chromosome) (Patterson and Stone 1949; Markow and O’Grady 2006; Vicoso and Bachtrog 2015). While the fusions of these elements differ between *Drosophila* species, the broad synteny is conserved. In our analysis we refer to the chromosomes by their Muller element names, instead of any species-specific chromosome name. Given the geographic distance between populations, we expected to find a corresponding signature of population differentiation across the populations for each Muller element. Using Structure (Falush *et al*. 2003), we find surprisingly little population differentiation between locations for each Muller element (Supplementary Figure 1C) but some structure by location for the mitochondrial genome (Supplementary Figure 1D), consistent with previous findings (Dyer 2004; Dyer and Jaenike 2005). Together these suggest that there is either still a large amount of standing ancestral variation shared between populations, or that there is gene flow between populations mostly via males.

Given that our populations are either recently established or have high levels of migration, we sought to determine which demographic model best explains the pattern of shared nuclear and mitochondrial variation seen in *D. innubila*. The relatively recent establishment of these populations suggests they may not be at migration-drift balance meaning that F_ST_ may not be an appropriate statistic to determine the extent of gene flow (Gutenkunst *et al*. 2010; Puzey *et al*. 2017). Using the silent nuclear polymorphism of each population in δaδi (Gutenkunst *et al*. 2010), we found the best fitting model for the nuclear variation in each pair of populations. In nearly all cases, the best model was one considering all populations as a single population following a population bottleneck and expansion, as opposed to separate populations with or without migration. This was true whether we considered all variation together or separately for each Muller element (Supplementary Table 3, *p*-value > 0.94). This suggests that the shared variation between populations is due to ancestrally maintained variation as opposed to elevated gene flow homogenizing new mutations arising in particular populations. Contrasting this, we find that the variation in mitochondrial genome fits a model of separate populations with some migration (mean migration rate = 0.0525, *p*-value < 0.00794) for all but one pair of populations (HU and SR best fit the model of a single population following a bottleneck, *p*-value = 0.423). This suggests that these populations are recently diverged, and HU and SR should be considered as a single population in all regards (Figure 1). The smaller effective population size of the mitochondrial genome may have resulted in it reaching migration-drift equilibrium faster than the nuclear genome.

Though not an appropriate model to determine population divergence, we used F_ST_ as a measure of shared variation (Weir and Cockerham 1984), and consistent with the δaδi models we find little differentiation between nuclear genomes (Figure 1B). We only find 65,899 nuclear SNPs (∼2% of called nuclear SNPs) are exclusive to populations, while most SNPs are shared between at least two populations (Supplementary Figure 2), suggesting most variation is maintained from before each population established separately. In contrast, there is higher F_ST_ between mitochondrial genomes in pairwise comparisons (Figure 1B). Both nuclear and mitochondrial δaδi and F_ST_ results are consistent with the Structure/StairwayPlot results (Supplementary Figure 1).

**Figure 2:**
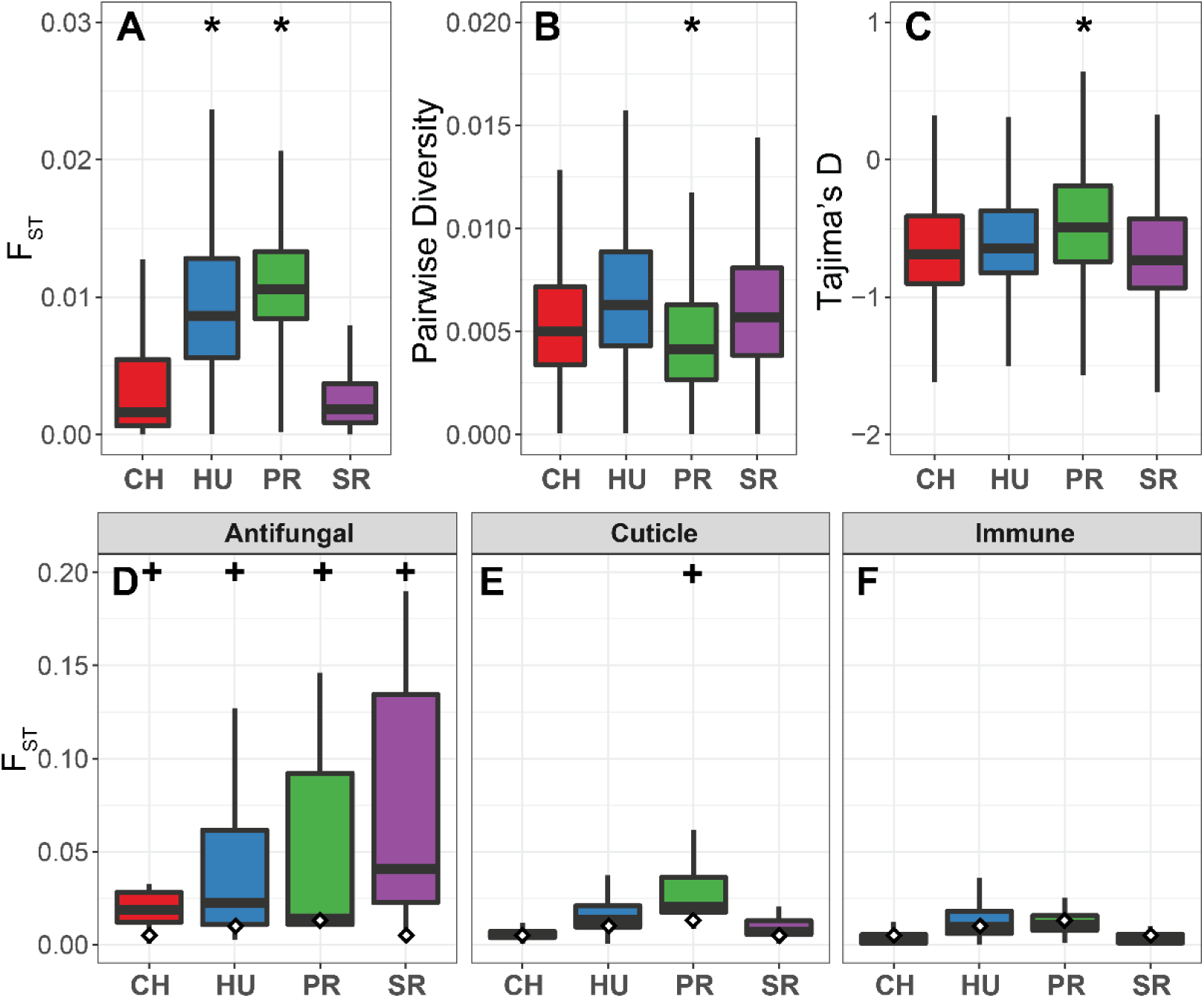
Summary statistics for each population. **A**. Distribution of F_ST_ per gene for each population versus all other populations. **B**. Distribution of genic pairwise diversity for each population. **C**. Distribution of genic Tajima’s D for each population. **D**. F_ST_ distribution for Antifungal associated genes for each population. **E**. F_ST_ distribution for cuticular proteins for each population. **F**. F_ST_ distribution for all immune genes (excluding antifungal genes). In A, B & C all cases significant differences from CH are marked with an * and outliers are removed for ease of visualization. In D, E & F, significant differences from the genome background in each population are marked with a + and white diamond mark the whole genome average of F_ST_ for each population.

We next determined whether the δaδi results were consistent with our expectations about a more recent expansion of *D. innubila* into Prescott. We find the time between population split for Prescott is significantly lower than all other comparisons (GLM t-value = -2.613, *p*-value = 0.0137), and the time since population expansion is also significantly lower for PR than other populations (GLM t-value = -2.313, *p*-value = 0.0275). F_ST_ of the nuclear genome is also significantly higher in Prescott comparisons (Figure 2A, GLM t-value = 93.728, *p*-value = 2.73e-102), though is still extremely low genome-wide in all pairwise comparisons involving PR (PR median = 0.0105), with some outliers on Muller element B like other populations (Supplementary Figures 3 & 4). We also calculated the population genetic statistics pairwise diversity and Tajima’s D for each gene using total polymorphism (Tajima 1989). As expected with a recent population contraction in Prescott (suggesting recent migration and establishment in a new location), pairwise diversity is significantly lower (Figure 2B, GLM t-value = -19.728, *p*-value = 2.33e-86, Supplementary Table 2) and Tajima’s D is significantly higher than all other populations (Figure 2C, GLM t-value = 4.39, *p*-value = 1.15e-05, Supplementary Table 2). This suggests that there is also a deficit of polymorphism in general in Prescott, consistent with a more recent population bottleneck, removing rare alleles from the population (Figure 2C, Supplementary Figure 5). Conversely, the other populations show a genome wide negative Tajima’s D, consistent with a recent demographic expansion (Supplementary Figure 5), suggesting a more recent bottleneck in Prescott relative to other populations.

**Figure 3:**
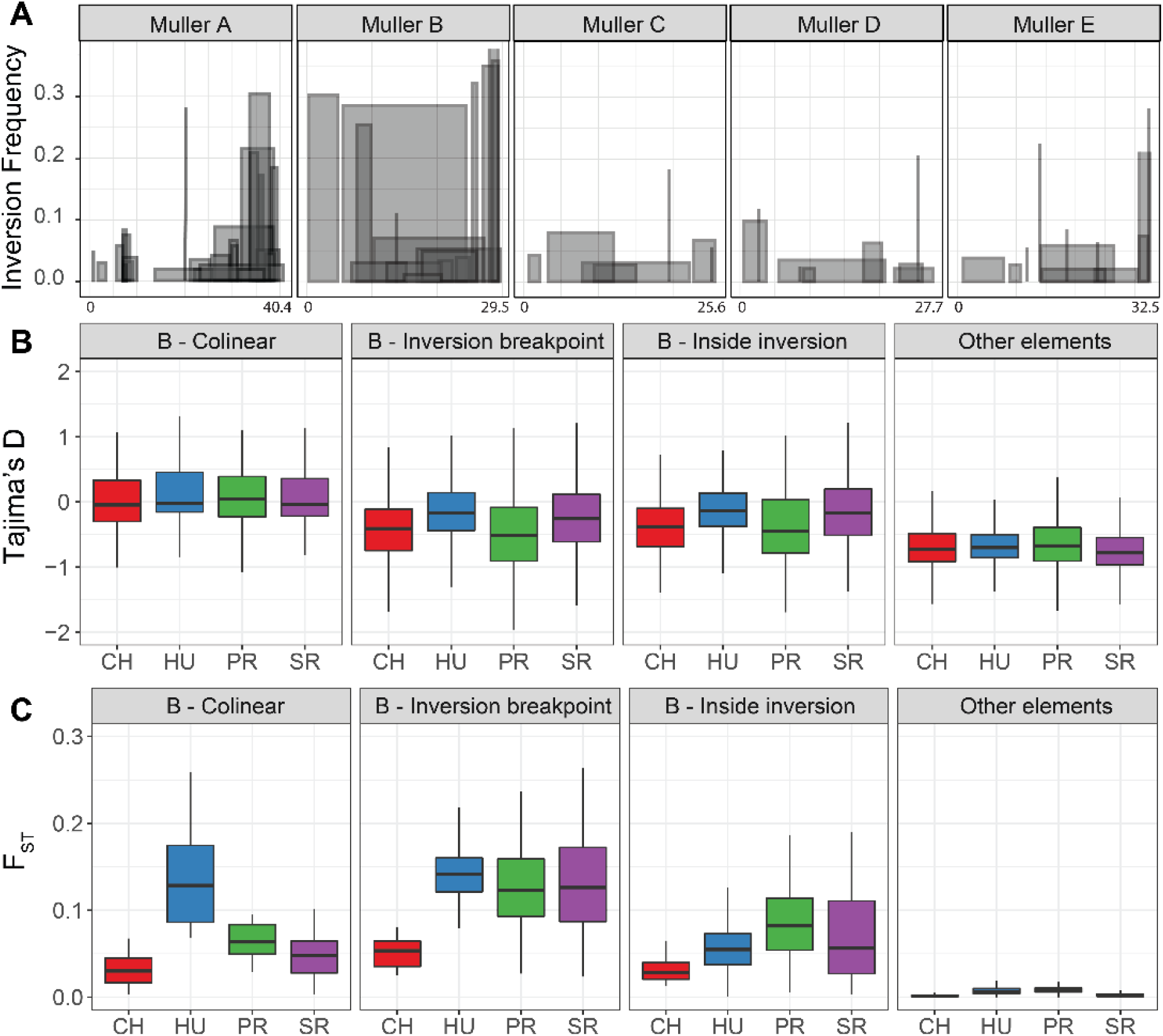
Summary of the inversions detected in the *Drosophila innubila* populations. **A**. Location and frequency in the total population of segregating inversions at higher than 1% frequency and greater than 100kbp. Shaded bars highlight the beginning and end of each inversion (X-axis) on each chromosome, as well as the inversion frequency (Y-axis). **B**. Tajima’s D and **C**. F_ST_ for genes across Muller element B, grouped by their presence under an inversion, outside of an inversion, near the inversion breakpoints (within 10kbp) or on a different Muller element. All inversions and frequencies inferred compared to the reference *D. innubila* genome.

**Figure 4:**
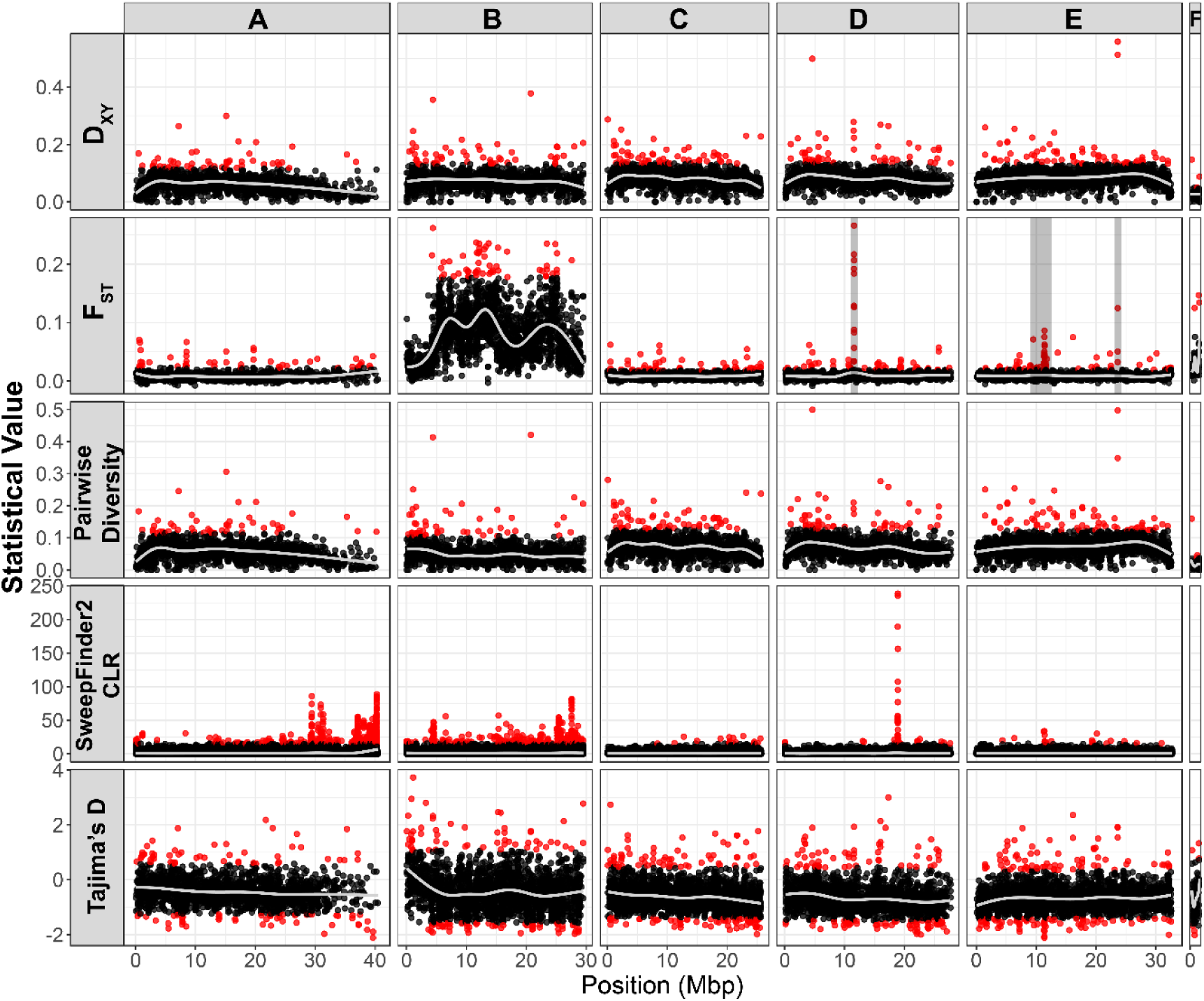
Comparison of estimated statistics across the *D. innubila* genome for the Prescott (PR) population. Values are as follows: the average pairwise divergence per gene (D_XY_, Prescott vs. all other populations), the gene wise population fixation index (F_ST_, Prescott vs. all other populations), within population pairwise diversity per genes, Composite Likelihood Ratio (CLR) per SNP calculated using Sweepfinder2, and gene wise within population Tajima’s D. For D_XY,_ F_ST_, Pairwise Diversity and Tajima’s D, Genes or windows in the upper 97.5^th^ percentile per chromosome are colored red. The lower 2.5^th^ percentile per chromosome is also colored in red for Tajima’s D. For Composite Likelihood Ratio the upper 99^th^ percentile windows are colored red. Shaded regions on the F_ST_ plot are further examined in Figure 5.

**Figure 5:**
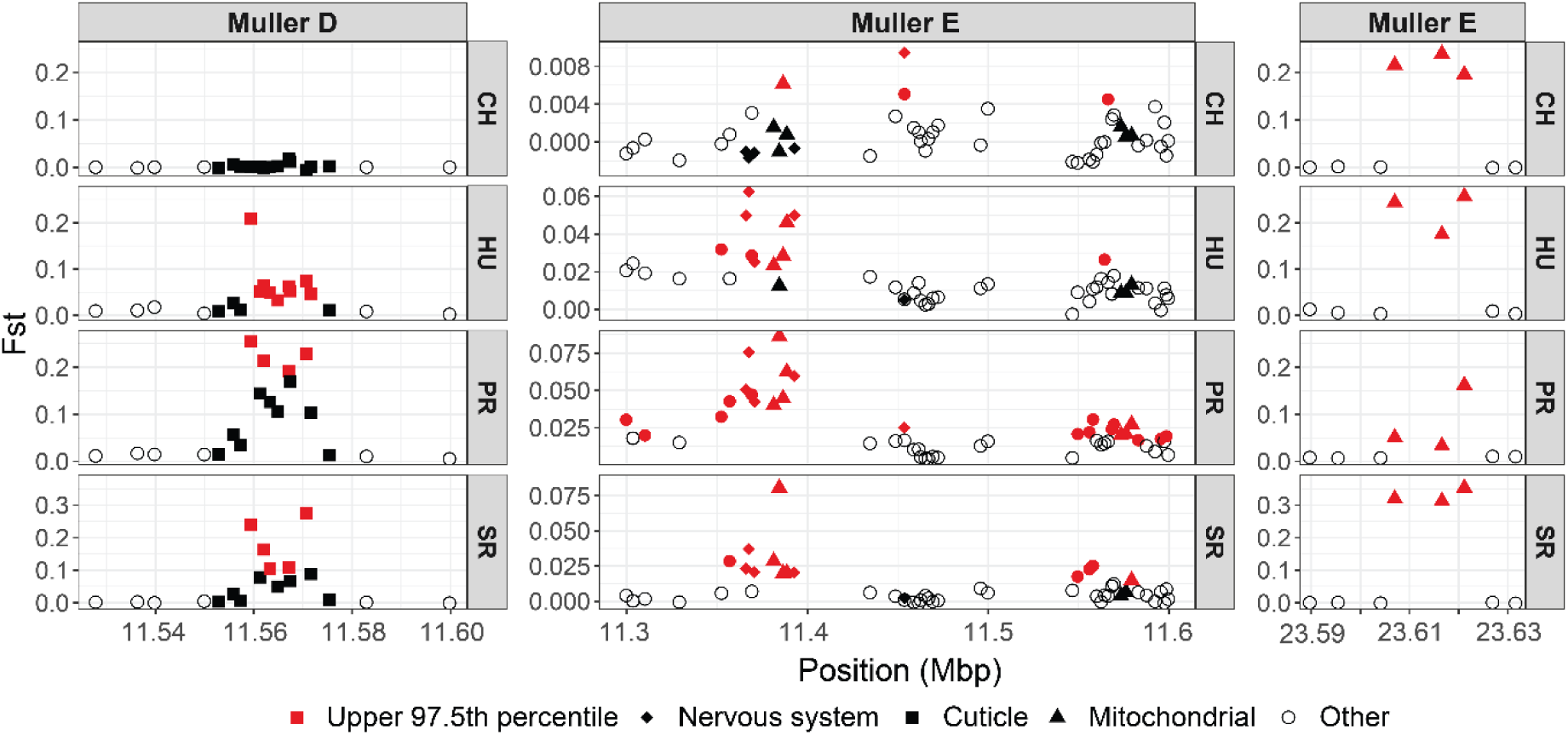
Gene-wise F_ST_ showing regions of elevated divergence between populations for each population. Plot shows F_ST_ for each gene in these regions to identify the causal genes. Genes with noted functions (cuticle development or mitochondrial translocations) are shown by point shape. Note the Y-axes are on different scales for each plot. Genes in the upper 97.5^th^ percentile of F_ST_ on each chromosome are colored in red.

### Population structure in the D. innubila genome is associated with segregating inversions

Though populations are panmictic for Muller element B (Using δaδi, Supplementary Table 3), F_ST_ is significantly higher on Muller element B compared to all other elements in all populations (Supplementary Figure 3, GLM t-value = 30.02, *p*-value = 3.567e-56) and is enriched for windows in the upper 97.5^th^ percentile of F_ST_ (Supplementary Figure 3, χ^2^ test: χ^2^ = 79.66, d.f. = 1, *p*-value = 1.06e-16). Additionally, we identified 10kbp windows across the genome in the upper 97.5^th^ percentile for the proportion of population-exclusive SNPs and found Muller B is significantly enriched for population-exclusive SNPs in every population (Supplementary Figure 3 & 4, χ^2^ test: χ^2^ = 60.28, d.f. = 1, *p*-value = 1.059e-12). All other chromosomes are significantly depleted for population-exclusive SNPs (χ^2^ test: χ^2^ = 31.90, d.f. = 1, *p*-value = 1.33e-6). On Muller element B, regions of elevated F_ST_ are consistent in all pairwise comparisons between populations (Supplementary Figure 4). Muller element B also has elevated Tajima’s D compared to all other Muller elements (Supplementary Figure 5, GLM t-value = 10.402, *p*-value = 2.579e-25), suggesting some form of structured population unique to Muller element B. However, actual values of F_ST_ are still low on Muller element B and populations are considered panmictic when fitting models with δaδi for all 1Mbp windows across Muller element B (Supplementary Table 3) (Gutenkunst *et al*. 2010), suggesting population structure by location is still minimal (Supplementary Figure 3, Supplementary Table 3, PR Muller B mean = 0.081, Muller C mean = 0.0087, Mitochondrial mean = 0.507).

We attempted to identify if this elevated structure is due to chromosomal inversions on Muller element B, comparing F_ST_ of a region to the presence or absence of inversions across windows (using only inversions called by both Delly and Pindel (Ye *et al*. 2009; Rausch *et al*. 2012)). We find several putative inversions across the genome at appreciable frequencies (89 total above 1% frequency), of which, 37 are found spread evenly across Muller element B and 22 are found at the telomeric end of Muller element A (Figure 3A). The presence of an inversion over a region of Muller element B is associated with higher genic F_ST_ in these regions (Figure 3A, Wilcoxon Rank Sum test W = 740510, *p*-value = 0.0129), though, when calculating F_ST_ for inversions, we find these inversions are not unique or even at different frequencies in specific populations (Mean inversion F_ST_ = 0.024, Max inversion F_ST_ = 0.22, χ^2^ test for enrichment in a specific population *p*-value > 0.361 for all inversions). Genes within 10kbp of an inversion breakpoint have significantly higher F_ST_ than outside the inverted regions, consistent with findings in other species (Figure 3C, GLM t-value = 7.702, *p*-value = 1.36e-14) (Machado *et al*. 2007; Noor *et al*. 2007). However, genes inside inverted regions show no difference in F_ST_ compared to those outside (Figure 3C, GLM t-value = -0.178, *p*-value = 0.859). All regions of Muller element B have higher F_ST_ than the other Muller elements (Figure 3C, outside inversions Muller element B vs all other chromosomes: GLM t-value = 7.379, *p*-value = 1.614e-13), suggesting some chromosome-wide force drives the higher F_ST_ and Tajima’s D, opposed to reduced recombination near inversion breakpoints (Noor *et al*. 2007).

Given that calls for large inversions in short read data are often not well supported (Chakraborty *et al*. 2017) and the apparently complex nature of the Muller element B inversions (Figure 3A), we may not have correctly identified the actual inversions and breakpoints on the chromosome. Despite this, our results do suggest a link between the presence of inversions on Muller element B and elevated differentiation of the entire Muller element B between *D. innubila* populations.

### Evidence for increased population differentiation in antifungal and cuticle development genes

Though differentiation is low across most of the genome in each population as populations are recently established and most variants are shared, we still find several genomic regions with relatively elevated differentiation, using F_ST_ as a proxy for divergence. In addition to the entirety of Muller element B, there are narrow windows of high F_ST_ on Muller elements D and E (Figure 4, Supplementary Figure 3 & 4). We attempted to identify whether any gene ontology groups have significantly higher F_ST_ than the rest of the genome. We consider a peak of elevated F_ST_ to be genes in the upper 97.5^th^ percentile of F_ST_ on each chromosome. These peaks for F_ST_ are enriched for antifungal genes in all populations and all in pairwise comparisons (Figure 2D, Supplementary Table 4, GO enrichment = 16.414, *p*-value = 1.61e-10). These antifungal genes (including *Dif, cactus* and *Trx-2* in all populations, *grass* in PR, and *modSP* in CH) are distributed across the genome and so not all under one peak of elevated F_ST._ Interestingly, this is the only immune category with elevated F_ST_ (Figure 2F), with most of the immune system showing no signatures of increased divergence between populations (Figure 2F, Supplementary Figure 6). This suggests that that fungal pathogens apply the strongest divergent selective pressure among populations.

**Figure 6:**
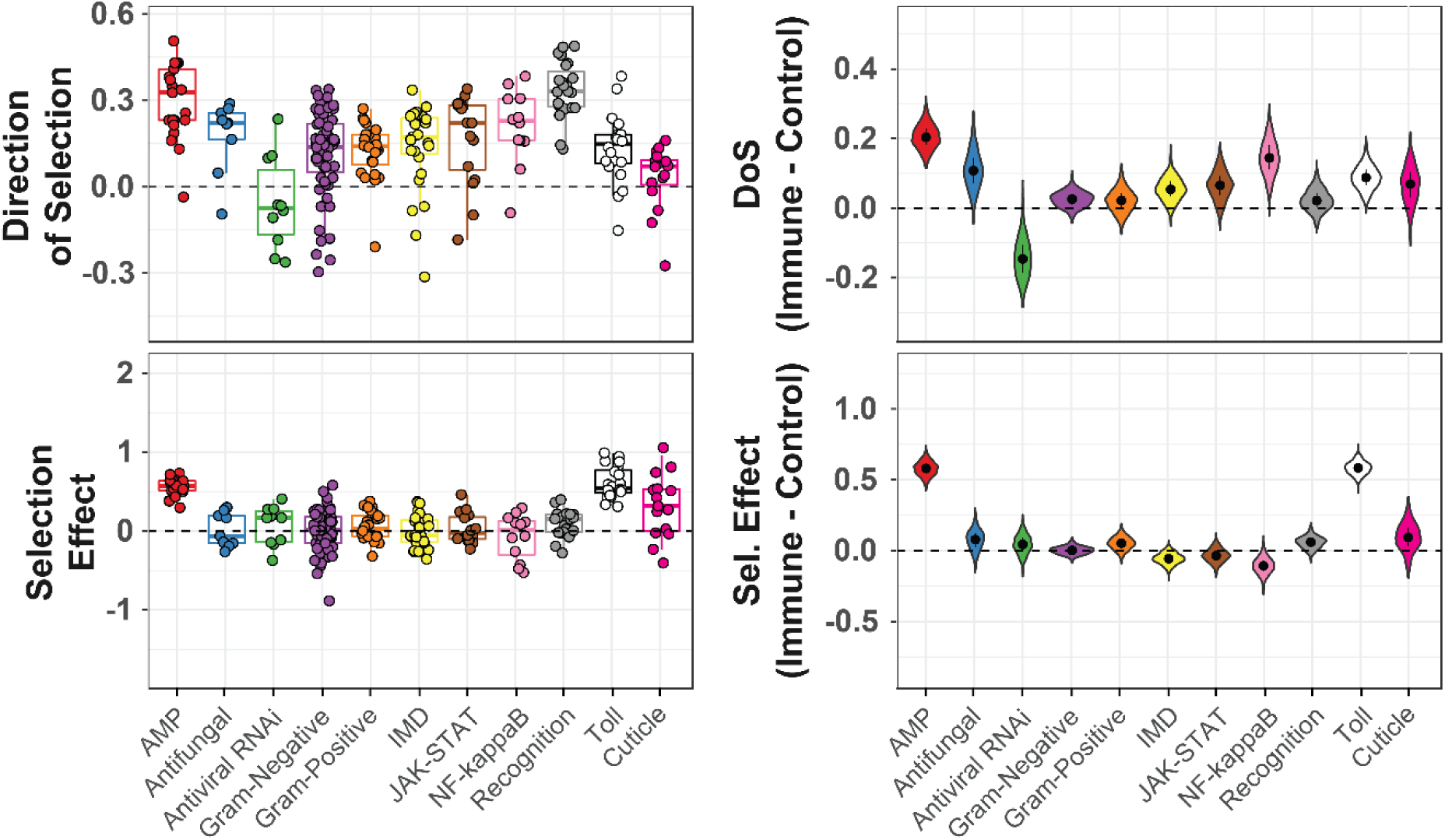
McDonald-Kreitman based statistics for immune categories in *D. innubila* and cuticle development. The left two plots show estimated statistics (Direction of Selection and Selection Effect) for each gene. The right two plots show the difference in mean statistic (Direction of Selection and Selection Effect) for each gene and a randomly sampled nearby gene.

Genes related to cuticle development also have significantly higher F_ST_, when compared to all other genes (Figure 2E, Supplementary Table 4, GO enrichment = 5.03, *p*-value = 8.68e-08), which could be associated with differences in the environment between locations (toxin exposure, humidity, *etc*.). Consistent with this result, the peak of F_ST_ on Muller element D (Figure 5, Muller element D, 11.56-11.58Mb) is composed exclusively of genes involved in cuticle development (e.g. *Cpr65Au, Cpr65Av, Lcp65Ad*) with elevated F_ST_ in these genes in all pairwise comparisons involved SR and HU, and the one vs all SR and HU population comparisons as well as PR comparisons (Figure 5), suggesting that they may be adapting to differing local conditions in those populations.

Two other clear peaks of elevated F_ST_ on Muller element E are also composed of genes in similar genomic categories (e.g. cuticle development). There also appears to be three regions of the *D. innubila* genome with translocated mitochondrial genes (Figure 5). The first peak (Muller element E, 11.35-11.4Mb) is composed exclusively of one of these translocated mitochondrial regions with 3 mitochondrial genes (including *cytochrome oxidase II*). The second peak (Muller element E, 23.60-23.62Mb) contains four other mitochondrial genes (including *cytochrome oxidase III* and *ND5*) as well as genes associated with nervous system activity (such as *Obp93a* and *Obp99c*). We find no correlation between coverage of these regions and mitochondrial copy number (Supplementary Table 1, Pearsons’ correlation t-value = 0.065, *p*-value = 0.861), so this elevated F_ST_ is probably not an artefact of mis-mapping reads. However, we do find these regions have elevated copy number compared to the rest of the genome (Supplementary Figure 7, GLM t-value = 9.245, *p*-value = 3.081e-20), and so this elevated divergence may be due to collapsed paralogs. These insertions of mtDNA are also found in *D. falleni* and are diverged from the mitochondrial genome, suggesting ancient transpositions (Hill *et al*. 2019). The nuclear insertions of mitochondrial genes are also enriched in the upper 97.5^th^ percentile for F_ST_ in HU and PR, when looking at only autosomal genes (Supplementary Table 4, GO enrichment = 4.53, *p*-value = 3.67e-04). Additionally, several other energy metabolism categories are in the upper 97.5^th^ percentile for F_ST_ for all pairwise comparisons involving CH. We find peaks of elevated pairwise diversity exclusively on the mitochondrial translocations (Supplementary Figure 8), suggesting unaccounted for variation in these genes which is consistent with duplications detected in these genes (Rastogi and Liberles 2005) (Supplementary Figure 7). This supports the possibility that unaccounted for duplications may be causing the elevated F_ST_ pairwise diversity in mitochondrial genes (Supplementary Figures 6-8). Overall these results suggest a potential divergence in the metabolic needs of each population, and that several mitochondrial genes may have found a new function in the *D. innubila* genome and may be diverging (and changing in copy number, Supplementary Figure 7) due to differences in local conditions.

**Figure 7:**
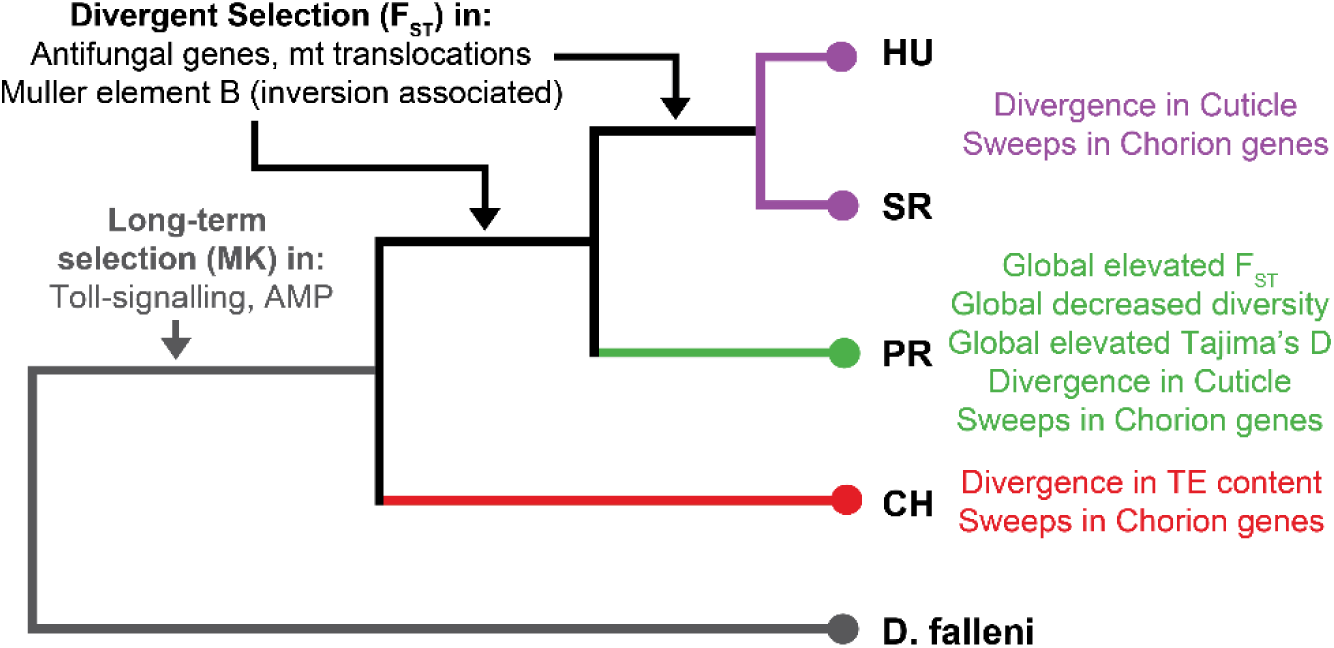
Summary of signatures of selection (Selective sweeps) and population differentiation (F_ST_ & D_XY_) shown on the phylogeny of *Drosophila innubila* populations. Signals seen as divergence between all populations (F_ST_) or recurrent/long-term evolution (McDonald-Kreitman, MK) is shown in black and grey respectively on the left-hand side. Signals seen on each branch are colored by the branches that the signals are seen on, shown on the right-hand side.

There has been considerable discussion over the last several years about the influence of demographic processes and background selection on inference of local adaptation (Cutter and Payseur 2013; Cruickshank and Hahn 2014; Hoban *et al*. 2016; Matthey-Doret and Whitlock 2018). In contrast to F_ST_ which is a relative measure of population differentiation, D_XY_ is an absolute measure that may be less sensitive to other population-level processes (Nei 1987; Cruickshank and Hahn 2014). In our data, the upper 97.5^th^ percentile for F_ST_ also significantly overlaps with the upper 97.5^th^ percentile for D_XY_ (Figure 4, Supplementary Figures 3, 4 & 8, χ^2^ text: χ^2^ = 62.61, p-value = 2.54e-14). The upper 97.5^th^ percentile for D_XY_ is enriched for chorion proteins in all pairwise comparisons and antifungal proteins for all pairwise comparisons involving PR (Supplementary Table 6, *p*-value < 0.05). Further, the high peak of F_ST_ over Muller element D cuticle proteins seen in all populations is also present when using D_XY_ (Figure 4). We find no evidence for duplications in the antifungal, cuticle or chorion proteins, suggesting the elevated F_ST_ and D_XY_ is likely due to local adaptation and not because of unaccounted for copy number variation (Figure 4, Supplementary Figure 7).

### Evidence for recent selective sweeps in chorion genes in each population

Recent adaptation often leaves a signature of a selective sweep with reduced polymorphism near the site of the selected variant. We attempted to identify selective sweeps in each population (and identify similar selective sweeps in multiple populations) using Sweepfinder2 to calculate the composite likelihood of a selective sweep (Huber *et al*. 2016). We first assessed if percentile of F_ST_ and composite likelihood overlap, suggesting that the elevated F_ST_ is due to a recent selective sweep (Nielsen 2005). We find no evidence of selective sweeps overlapping with genes with elevated F_ST_ (Supplementary Figure 9A, χ^2^ test for overlap of 97.5^th^ percentile windows χ^2^ = 1.33 *p*-value = 0.249), suggesting that the divergence between populations is likely not because of local adaptation. This is consistent with the high levels of ancestrally maintained variation we find and the recent establishment of each population (Figure 1). The upper 97.5^th^ percentile of composite likelihood is equally distributed across the genome and are not enriched for any gene ontology categories (*p-*value > 0.31), but if we limit our survey, we find the upper 99^th^ percentile is enriched for chorion/egg development genes in all populations (Enrichment = 5.12, *p*-value = 0.00351).

We find one peak of extremely high composite likelihood values present in all populations on Muller element D, but most extreme in Prescott (Figure 4, Supplementary Figure 9B, Muller D, 18.75-19Mb). The center of this peak is upstream of the cuticle protein *Cpr66D* and four chorion proteins (*Cp15, Cp16, Cp18, Cp19*), with several cell organization proteins (*Zasp66, Pex7, hairy, Prm, Fhos*) within 10kbp of the sweep center. These specific chorion proteins do not have elevated F_ST_ or D_XY_ compared to the rest of the genome (Wilcoxon Rank Sum W = 399170, *p*-value = 0.273), but has significantly elevated D_XY_ compared to other genes within 50kbp (Supplementary Figures 8 & 9, Wilcoxon Rank Sum W = 45637000, *p*-value = 0.0158). However, this may be due to the reduced diversity seen in these chorion genes because of the selective sweep (McVean 2007).

Finally, we find evidence of several peaks of selective sweeps in the non-recombining telomere of the X chromosome (Muller A, 39.5-40.5Mb), among several uncharacterized genes (Supplementary Figure 9). Consistent with this, we find F_ST_ is elevated between our 2001 and 2017 Chiricahua samples, suggesting at least one recent sweep on the X telomere (Supplementary Figure 10A), resulting in different variants fixing in the 16 years between samplings.

### Toll-related immune show signatures of recurrent positive selection in D. innubila

Finally, we sought to if identify genes and functional categories showing strong signatures of selective sweeps and divergence are also undergoing adaptive evolution, suggesting species-wide long-term evolution as opposed to recent adaptation. We reasoned that if the elevated population differentiation seen in antifungal genes and cuticle development proteins (Figure 2 & 3, Supplementary Figure 6) was due to adaptation also acting over longer time periods, we would expect to see signatures of ancient adaptation in those categories. Furthermore, Hill *et al*. used d_N_/d_S_-based statistics to show that genes involved in some immune defense pathways were among the fastest evolving genes in the *D. innubila* genome (Hill *et al*. 2019). We also sought to identify what genes are evolving due to recurrent positive selection in *D. innubila* in one or all populations. To this end we calculated the McDonald-Kreitman based statistic direction of selection (DoS) (Stoletzki and Eyre-Walker, 2011) and SnIPRE selection effect (Eilertson et al., 2012) to estimate the proportion of substitutions fixed by adaptation and identify genes under recurrent selection. We then fit a linear model to identify gene ontology groups with significantly higher DoS or selection effect than expected. In this survey we found cuticle genes and antifungal genes did have some signatures of adaptive evolution (DoS > 0, Selection Effect > 0 and significant McDonald-Krietman test for 80% of genes in these categories) but as a group showed no significant differences from the background (GLM t-value = 1.128, *p*-value = 0.259, Supplementary Table 5). In fact, we only found two functional groups significantly higher than the background, Toll signaling proteins (GLM t-value = 2.581 *p*-value = 0.00986, Supplementary Table 4) and antimicrobial/immune peptides (AMPs, GLM t-value = 3.66 *p*-value = 0.00025, Supplementary Table 4). In a previous survey we found that these categories also had significantly elevated rates of amino acid divergence (Hill *et al*. 2019). These results suggest that this divergence is indeed adaptive. We find no evidence that genes under recent selective sweeps are undergoing recurrent evolution, suggesting these genes are only recent targets of selection (Supplementary Table 4).

Five immune peptides showed consistently positive DoS and selection effect values (which are also among the highest in the genome): four *Bomanins* and *Listericin*. The *Bomanins* are immune peptides regulated by Toll signaling, while JAK-STAT regulates *Listericin* (Hoffmann 2003; Takeda and Akira 2005). *Listericin* has been implicated in the response to viral infection due to its expression upon viral infection (Dostert *et al*. 2005; Zambon *et al*. 2005; Imler and Elftherianos 2009; Merkling and van Rij 2013), additionally Toll-regulated AMPS have been shown to interact with the dsDNA Kallithea virus in *Drosophila melanogaster* (Palmer et al. 2018). *D. innubila* is burdened by Drosophila innubila nudivirus (DiNV), a Nudivirus closely related to Kallithea virus, that infects 40-80% of individuals in the wild (Unckless 2011; Hill and Unckless 2020). Consistent with the adaptation observed in Toll signaling proteins, this suggests the Toll immune system is under long term selection to resist infection by this DNA virus. However, Toll is also the major component of humoral defense against Gram-positive bacteria and fungi (Kimbrell and Beutler 2001). Since *D. innubila* feed and breed in rotting mushrooms (Jaenike and Perlman 2002), we can’t rule out that the recurrent adaptation in Toll genes is related to adaptation to the conditions of their pathogen-infested habitats. For all immune categories, as well as cuticle proteins and antifungal proteins, we find no significant differences between populations for either MK-based statistics, and no significant differences in the distribution of these statistics between populations (GLM t-value < 0.211, p-value > 0.34 for all populations, Supplementary Table 4). Selection at these loci is likely long term due to an ancient selective pressure and has been occurring since before the separation of these populations. Mutation rates, efficacy of selection and population structure can vary across the genome, which can confound scans for selection (Charlesworth *et al*. 2003; Stajich and Hahn 2005).To work around this, we employed a control-gene resampling approach to identify the average difference from the background for each immune category (Chapman *et al*. 2019). We find no signatures of ancient positive selection in antifungal genes (Supplementary Figure 11, 61% resamples > 0) or cuticle genes (Figure 6, 54% resamples > 0) but do again find extremely high levels of positive selection in AMPs (Figure 6, 100% resamples > 0) and Toll signaling genes (Figure 6, 99.1% resamples > 0). Segregating slightly deleterious mutations can bias inference of selection using McDonald-Kreitman based tests (Messer and Petrov 2012). To further account for this bias, we also calculated asymptotic α for all functional categories across the genome (Haller and Messer 2017). As before, while we find signals for adaptation in antifungal and cuticle proteins (asymptotic α > 0), values in both categories do not differ from the background (Supplementary Figure 11, permutation test Antifungal *p*-value = 0.243, Cuticle *p*-value = 0.137). As with the basic McDonald-Kreitman statistics (Figure 6), the only categories significantly higher than the background are Toll signaling genes (Permutation test *p*-value = 0.033) and AMPs (Permutation test *p*-value = 0.035). Together these results suggest that while genes involved in antifungal resistance and cuticle development are slightly divergent between populations (Figure 5), this evolution has likely not been occurring over long-time scales and so is restricted to their current environments. Alternatively, the adaptation may be too recent to detect a signal using polymorphism and divergence-based metrics. In either case, signals of longer-term adaptation in *D. innubila* appear to be driven by host-pathogen interactions (possibly with the long-term viral infection DiNV (Hill *et al*. 2019)) as opposed to local adaptation.

## Discussion

Interpopulation divergence is a time-dependent process where pro-divergence factors of genetic drift and local adaptation oppose ancestral variation and migration (Rankin and Burchsted 1992; Gillespie 2004; Excoffier *et al*. 2009; White *et al*. 2013). We sought to examine the extent that these factors drive divergence, and the genomic basis of colonization, divergence and local adaptation in *Drosophila innubila*, a geographically structured species found across four forests separated by large expanses of desert. We characterized the phylogeographic history of four populations of *Drosophila innubila* (Figure 7), a mycophagous species endemic to the Arizonan Sky islands using whole genome resequencing of 318 wild-caught individuals. *D. innubila* expanded into its current range during or following the previous glacial maximum (Figure 1, Supplementary Figure 1). Interestingly, there is very little support for population structure across the nuclear genome (Figures 1 & 2, Supplementary Figures 1-4), including in the repetitive content (Supplementary Figure 12), but some evidence of population structure in the mitochondria, as found previously in *D. innubila* (Dyer and Jaenike 2005). There are two models which could explain this pattern: the first is extremely high migration, but of mostly males (as seen in other disparate species (Rankin and Burchsted 1992; Searle *et al*. 2009; Ma *et al*. 2013; Avgar and Fryxell 2014)), resulting in a panmictic nuclear genome but diverged mitochondrial genome (as shown in Figure 1B). The second model (which we found better support for using δaδi, Supplementary Table 3) is that ancestral variation is maintained due to the recent isolation of populations (less than 4N_e_ generations ago) and recent coalescent times (Figure 1B, Supplementary Figure 3), though the elevated mutation rate and reduced N_e_ of the mitochondria results in an increased rate of differentiation by drift between mitochondrial genomes (Gillespie 2004). Similarly, Drosophila innubila Nudivirus, a DNA virus infecting *Drosophila innubila*, is also highly structured, possibly due to the smaller N_e_, elevated mutation rate and high levels of adaptation (Hill and Unckless 2020). *D. innubila* is also parasitized by a male-killing strain of the maternally transmitted, intracellular bacterium, *Wolbachia*. This alphaproteobacterium infects about 65% of arthropods (Werren *et al*. 2008), and has evolved several strategies to improve their chances of transmission between generations, including killing male offspring in *D. innubila* (Dyer and Jaenike 2005; Dyer *et al*. 2005; Jaenike and Dyer 2008; Werren *et al*. 2008). The mitochondrial divergence could be driven by linkage to the *Wolbachia*: when *Wolbachia* variants sweep (such as variants that improve parasitism), mitochondrial variants will hitchhike to fixation (Jaenike *et al*. 2003; Dyer and Jaenike 2005). This will then further decrease the effective population size of the mitochondria, reaching migration-drift equilibrium faster than expected for a haploid genome under drift.

Most Sky Island species are highly diverged between locations, suggesting that *D. innubila* may be much more recently established than other species (Smith and Farrell 2005; Fave *et al*. 2015; Wiens *et al*. 2019). Similar patterns of limited nuclear divergence between species are observed in a Sky Island bird species, *S. carolinensis* (Manthey and Moyle 2015), however this flying species can easily migrate between Sky Islands (Manthey and Moyle 2015). In contrast, the patterns observed in *D. innubila* populations are more likely ancestrally maintained and not due to gene flow (Figure 1 & 2, Supplementary Figure 2). In *S. carolinensis*, differential selection is restricted to loci associated with recent changes in environmental extremes (Manthey and Moyle 2015). This may also be the case in *D. innubila*, as we find some evidence of recent selection, limited to genes associated with environmental factors, chorion genes and antifungal immune genes. (Figure 7, Supplementary Figure 2). Most other insect and non-flying species found across the Sky Islands are highly structured between locations (Smith and Farrell 2005; Fave *et al*. 2015; Wiens *et al*. 2019). If the lack of geographic divergence is due to the maintenance of ancestral variation, we expect *D. innubila* populations to become as structured as these other species over time.

Segregating inversions are often associated with population structure (Dobzhansky and Sturtevant 1937; Fuller *et al*. 2016; Puzey *et al*. 2017) and could explain the excess interpopulation divergence seen on Muller element B here (Supplementary Figures 2-5). Our detection of several putative segregating inversions on Muller element B relative to all other chromosomes (Figure 3A) supports this assertion. However, all are large and common inversions characterized in all populations, suggesting the inversions are not driving the elevated F_ST_. We suspect that the actual causal inversions may not have been correctly characterized for two reasons. First there are limitations of detecting inversions in repetitive regions with short read data (Marzo *et al*. 2008; Chakraborty *et al*. 2017). Second, and confounding the first, the likely existence of several overlapping segregating inversions complicates correctly calling inversion breakpoints further. The elevated F_ST_ could also be caused by other factors, such as extensive duplication and divergence on Muller element B being misanalysed as just divergence. In fact, the broken and split read pairs used to detect inversions are very similar to the signal used to detect duplications (Ye *et al*. 2009; Rausch *et al*. 2012; Chen *et al*. 2016), suggesting some misidentification may have occurred. If a large proportion of Muller B was duplicated, we would see elevated mean coverage of Muller element B in all strains compared to other autosomes, which is not the case (Supplementary Table 1). There may also be seasonal variation in inversions which causes the elevated Tajima’s D and F_ST_ seen here (Supplementary Figures 3-5) and may be the cause of the difference in minor allele frequency between 2001 and 2017 on Muller element B (Supplementary Figure 10B). In fact, in several species, inversions can change in frequency as the multiple linked variants they contain change in fitness over time (Rodriguez-Trelles *et al*. 1996; Oneal *et al*. 2014; Kapun *et al*. 2016; Puzey *et al*. 2017).

While we find no differences in inversion frequencies between time points (2001 and 2017) in *D. innubila* (Wilcoxon Rank Sum Test W = 561, *p*-value = 0.352), we again may not have correctly identified the inversions and so cannot properly infer frequency changes. Segregating inversions, such as those on Muller element B, are frequently associated with local or temporal variation (Dobzhansky and Sturtevant 1937; Rodriguez-Trelles *et al*. 1996; Oneal *et al*. 2014), and consistent with this find differences in allele frequencies between time points (Supplementary Figure 10B). There was an extensive forest fire in the Sky Islands in 2011 which could plausibly have been a strong selective force driving a change in allele frequency on Muller B between time points (Supplementary Figure 10B) (Arechederra-Romero 2012). It is also worth noting that *D. pseudoobscura* segregates for inversions on Muller element C and these segregate by population in the same Sky island populations (and beyond) as the populations described here (Dobzhansky and Sturtevant 1937; Dobzhansky *et al*. 1963; Fuller *et al*. 2016). The inversion polymorphism among populations is a plausible area for local adaptation and may provide an interesting contrast to the well-studied *D. pseudoobscura* inversions. (Hermisson and Pennings 2005; McVean 2007; Przeworski *et al*. 2011).

Ours is one of few studies that sequences whole genomes from individual wild-caught *Drosophila* and therefore avoids several generations of inbreeding that would purge recessive deleterious alleles (Gillespie 2004; Mackay *et al*. 2012; Pool *et al*. 2012). During inbreeding (or due to unintentional inbreeding over generations as lines are maintained) a large proportion of variation found in the population will be removed by background selection (Charlesworth *et al*. 1993). This limits the scope of studies, and the ability to infer the history of the population. Additionally, when attempting to calculate the extent of adaptation occurring in populations, inbred panels will have removed a large proportion of segregating (likely deleterious) variation, which may lead to misestimations of McDonald-Kreitman based statistics, which normalize divergence to polymorphism (McDonald and Kreitman 1991a; Eilertson *ET AL*. 2012; Messer and Petrov 2012). This could also explain the excessive rates of adaptation in *Drosophila melanogaster* compared to other species (Cai *et al*. 2009; Sella *et al*. 2009). The excess of putatively deleterious alleles which may be present in our study harkens back to early studies of segregating lethal mutations in populations as well as recent work on humans (Dobzhansky *et al*. 1963; Marinkovic 1967; Dobzhansky and Spassky 1968; Watanabe *et al*. 1974; Gao *et al*. 2015).

To date, most of the genomic work concerning the phylogeographic distribution and dispersal of different *Drosophila* species has been limited to the *melanogaster* supergroup (Pool *et al*. 2012; Pool and Langley 2013; Behrman *et al*. 2015; Lack *et al*. 2015; Machado *et al*. 2015), with some work in other *Sophophora* species (Fuller *et al*. 2016). This limits our understanding of how non-commensal species disperse and behave, and what factors seem to drive population demography over time. Here we have glimpsed into the dispersal and history of a species of mycophageous *Drosophila* and found evidence of changes in population distributions potentially due to the changing climate (Figure 7) (Arizona-Geological-Survey 2005) and population structure possibly driven by segregating inversions. Because many species have recently undergone range changes or expansions (Excoffier *et al*. 2009; Porretta *et al*. 2012; White *et al*. 2013), we believe examining how this has affected genomic variation is important for population modelling and even for future conservation efforts (Excoffier *et al*. 2009; Coe *et al*. 2012).

## Methods

### Fly collection, DNA isolation and sequencing

We collected wild *Drosophila* at the four mountainous locations across Arizona between the 22^nd^ of August and the 11^th^ of September 2017: the Southwest research station in the Chiricahua mountains (CH, ∼5,400 feet elevation, 31.871 latitude -109.237 longitude, 96 flies), in Prescott National Forest (PR, ∼7,900 feet elevation, 34.586 latitude -112.559 longitude, 96 flies), Madera Canyon in the Santa Rita mountains (SR, ∼4,900 feet elevation, 31.729 latitude -110.881 longitude, 96 flies) and Miller Peak in the Huachuca mountains (HU, ∼5,900 feet elevation, 31.632 latitude -110.340 longitude, 53 flies) (Coe *et al*. 2012). *Drosophila innubila* only emerge for a short period of time in the Arizona wet season (∼2 months), limiting dates of collection to this period of time each year (Patterson and Stone 1949). Baits consisted of store-bought white button mushrooms (*Agaricus bisporus*) placed in large piles about 30cm in diameter, with at least 5 baits per location. We used a sweep net to collect flies over the baits in either the early morning or late afternoon between one and three days after the bait was set. We sorted flies by sex and species at the University of Arizona in Tucson, AZ and flash frozen at -80°C before shipping on dry ice to the University of Kansas in Lawrence KS.

We sorted 343 flies (172 females and 171 males) which phenotypically matched *D. innubila*. We then homogenized and extracted DNA using the Qiagen Gentra Puregene Tissue kit (USA Qiagen Inc., Germantown, MD, USA), isolating DNA for each fly individually. We also isolated the DNA of 40 *D. innubila* samples individually, collected in early September in 2001 from CH. We prepared a genomic DNA library of these 383 DNA samples using a modified version of the Nextera DNA library prep kit (∼ 350bp insert size) meant to conserve reagents, during this process we barcoded each sample to distinguish individuals during sequencing. We sequenced the libraries on four lanes of an Illumina HiSeq 4000 (150bp paired end) (Supplementary Table 1, Data deposited in the NCBI SRA: SRP187240).

### Sample filtering, mapping and alignment

We removed adapter sequences using Scythe (Buffalo 2018), trimmed all data using cutadapt to remove barcodes (Martin 2011) and removed low quality sequences using Sickle (parameters: -t sanger -q 20 -l 50) (Joshi and Fass 2011). We masked the *D. innubila* reference genome, using *D. innubila* TE sequences generated previously and RepeatMasker version 4.0 (parameters: -s -gccalc -gff -lib customLibrary) (Smit and Hubley 2013-2015; Hill *et al*. 2019). We then mapped the short reads to the masked *D. innubila* genome using BWA MEM Version 0.7.17 (Li and Durbin 2009), and sorted and indexed using SAMTools Version 1.9 (Li *et al*. 2009). Following mapping, we added read groups, marked and removed sequencing and optical duplicates, and realigned around indels in each mapped BAM file using Picard Version 2.23.4 and GATK Version 4.0.0.0 (Http://broadinstitute.github.io/picard ; McKenna *et al*. 2010; DePristo *et al*. 2011). We then removed individuals with low coverage of the *D. innubila* genome (less than 5x coverage for 80% of the non-repetitive genome), and individuals we suspected of being misidentified as *D. innubila* following collection due to anomalous mapping. This left us with 280 *D. innubila* wild flies (48 - 84 flies per populations) from 2017 and 38 wild flies from 2001 with at least 5x coverage across at least 80% of the euchromatic genome (Supplementary Table 1).

### Nucleotide polymorphisms across the population samples

For the 318 sequenced samples with reasonable coverage, we called SNPs using GATK HaplotypeCaller Version bcf4.0.0.0 (McKenna *et al*. 2010; DePristo *et al*. 2011) which generated a multiple strain VCF file. We then used BCFtools Version 1.7 (Narasimhan *et al*. 2016) to remove sites with a GATK quality score (a composite PHRED score for multiple samples per site) lower than 950 and sites absent (e.g. sites of low quality, or with 0 coverage) from over 5% of individuals. This filtering left us with 4,522,699 SNPs and small indels across the 168Mbp genome of *D. innubila*. We then removed SNPs found as a singleton in a single population (as possible errors), leaving us with 3,240,198 SNPs. We used the annotation of *D. innubila* and SNPeff Version 5.0 (Cingolani *et al*. 2012) to identify SNPs as synonymous, non-synonymous, non-coding or another annotation. Simultaneous to the *D. innubila* population samples, we also mapped genomic information from outgroup species *D. falleni* (SRA: SRR8651761) and *D. phalerata* (SRA: SRR8651760) to the *D. innubila* genome and called divergence using the GATK variation calling pipeline to identify derived polymorphisms and fixed differences in *D. innubila*.

### Population genetic summary statistics and structure

Using the generated total VCF file with SNPeff annotations, we created a second VCF containing only synonymous polymorphism using BCFtools Version 1.7 (Narasimhan *et al*. 2016). We calculated pairwise diversity per base, Watterson’s theta, Tajima’s D (Tajima 1989) and F_ST_ (Weir and Cockerham 1984) (versus all other populations) across the genome for each gene in each population using VCFtools Version 0.1.13 (Danecek *et al*. 2011) and the VCF containing all variants. Using ANGSD Version 0.9.11 to parse the synonymous polymorphism VCF (Korneliussen *et al*. 2014), we generated synonymous unfolded site frequency spectra for the *D. innubila* autosomes for each population, using the *D. falleni* and *D. phalerata* genomes as outgroups to the *D. innubila* genome (Hill *et al*. 2019).

We used the population silent SFS with previously estimated mutation rates of *Drosophila* (Schrider et al. 2013), as inputs in StairwayPlot Version 2 (Liu and Fu 2015), to estimate the effective population size backwards in time for each location.

We also estimated the extent of population structure across samples using Structure Version 2.3.4 (Falush *et al*. 2003), repeating the population assignment for each chromosome separately using only silent polymorphism, for between one and ten populations (k = 1-10, 100000 iterations burn-in, 400000 iterations sampling). Following (Frichot *et al*. 2014), we manually assessed which number of subpopulations best fits the data for each *D. innubila* chromosome and DiNV to minimize entropy.

As our populations have recently established, we used δaδi Version 1.6.7 to determine which model of population dynamics best fits for each *D. innubila* chromosome (Gutenkunst *et al*. 2010).For pairs of populations, we compared models to determine if populations are behaving as one population having gone through a bottleneck (*bottlegrowth* model), as split populations with migration (*bottlegrowth_split_mig* model) or recently split populations with no migration (*bottlegrowth_split* model). We generated a site frequency spectrum for each chromosome for each population from our VCF of silent polymorphism in δaδi (though using the total polymorphism for the mitochondria), using the *Spectrum*.*from_data_dict* function. For each pair of populations and each chromosome, we estimated the optimal parameters for each model using the *Inference*.*optimize_log* function. We then fit each model (using the *Inference*.*ll_multinom* function) and compared the fit of each model using a likelihood-ratio test, to determine what demographic model best fits each Muller element and the mitochondria.

### Signatures of local adaptive divergence across D. innubila populations

We downloaded gene ontology groups from Flybase (Gramates *et al*. 2017). We then used a gene enrichment analysis to identify enrichments for particular gene categories among genes in the upper and lower 2.5^th^ percentile for F_ST_, Tajima’s D and Pairwise Diversity versus all other genes (Subramanian *et al*. 2005). Due to differences on the chromosomes Muller A and B versus other chromosomes in some cases, we also repeated this analysis chromosome by chromosome, taking the upper 97.5^th^ percentile of each chromosome.

We next attempted to look for selective sweeps in each population using Sweepfinder2 (Huber *et al*. 2016). We reformatted the polarized VCF file to a folded allele frequency file, showing allele counts for each base. We then used Sweepfinder2 Version 1.0 on the total called polymorphism in each population to detect selective sweeps in 1kbp windows (Huber *et al*. 2016). We reformatted the results and looked for genes neighboring or overlapping with regions where selective sweeps have occurred with a high confidence, shown as peaks above the genomic background. We surveyed for peaks by identifying 1kbp windows in the 97.5^th^ percentile and 99^th^ percentile for composite likelihood ratio per chromosome.

Using the total VCF with outgroup information, we next calculated D_XY_ per SNP for all pairwise population comparisons (Nei and Miller 1990), as well as within population pairwise diversity and d_S_ from the outgroups, using a custom python script. We then found the average D_XY_ and d_S_ per gene and looked for gene enrichments in the upper 97.5^th^ percentile, versus all other genes.

### Inversions

For each sample, we used Delly Version 2.0 (Rausch *et al*. 2012) to generate a multiple sample VCF file identifying regions in the genome which are potentially duplicated, deleted or inverted compared to the reference genome. Then we filtered and removed inversions found in fewer than 1% of individuals and with a GATK VCF quality score lower than 200. We also called inversions using Pindel Version 0.2.5 (Ye *et al*. 2009) in these same samples and again removed low quality inversion calls. We next manually filtered samples and merged inversions with breakpoints within 1000bp at both ends and significantly overlapping in the presence/absence of these inversions across strains (using a χ^2^ test, *p*-value < 0.05). We also filtered and removed large inversions which were only found with one of the two tools. Using the remaining filtered and merged inversions we estimated the frequency of each inversion within the total population.

### Signatures of recurrent selection

We filtered the total VCF with annotations by SNPeff and retained only non-synonymous (replacement) or synonymous (silent) SNPs (Cingolani *et al*. 2012). We then compared these polymorphisms to the differences identified to *D. falleni* and *D. phalerata* to polarize changes to specific branches. Specifically, we sought to determine sites which are polymorphic in our *D. innubila* populations or are substitutions which fixed along the *D. innubila* branch of the phylogeny. We used the counts of fixed and polymorphic silent and replacement sites per gene to estimate McDonald-Kreitman-based statistics, specifically direction of selection (DoS) (McDonald and Kreitman 1991b; Smith and Eyre-Walker 2002; Stoletzki and Eyre-Walker 2011). We also used these values in SnIPRE (Eilertson *et al*. 2012), which reframes McDonald-Kreitman based statistics as a linear model, taking into account the total number of non-synonymous and synonymous mutations occurring in user defined categories to predict the expected number of these substitutions and calculate a selection effect relative to the observed and expected number of mutations (Eilertson *et al*. 2012). We calculated the SnIPRE selection effect for each gene using the total number of mutations on the chromosome of the focal gene. Using FlyBase gene ontologies (Gramates *et al*. 2017), we sorted each gene into a category of immune gene or classed it as a background gene, allowing a gene to be classed in multiple immune categories. We fit a GLM to identify functional categories with excessively high estimates of adaptation, considering multiple covariates:

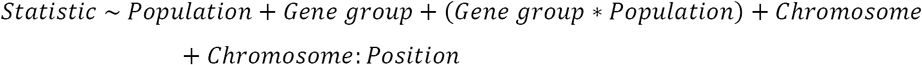

We then calculated the difference in each statistic between our focal immune genes and a randomly sampled nearby (within 100kbp) background gene, finding the average of these differences for each immune category over 10000 replicates, based on (Chapman *et al*. 2019).

To confirm these results, we also used AsymptoticMK (Haller and Messer 2017) to calculate asymptotic α for each gene category. We generated the non-synonymous and synonymous site frequency spectrum for each gene category, which we then used in AsymptoticMK to calculate asymptotic α and a 95% confidence interval. We then used a permutation test to assess if functional categories of interest showed a significant difference in asymptotic α from the rest of categories.

## Acknowledgements

This work was completed with helpful discussion from Justin Blumensteil, Joanne Chapman, Richard Glor, Stuart MacDonald, Maria Orive and Caroyln Wessinger. We would especially like to thank Kelly Dyer and Paul Guinsberg for proposing the idea of the manuscript and providing feedback on early sections of the manuscript. We greatly appreciate help provided by John Kelly, for providing scripts to calculate D_XY_ as well as advice on population genetic inference and comments on the manuscript. Collections were completed with assistance from Todd Schlenke and the Southwest Research Station. We thank Brittny Smith and the KU CMADP Genome Sequencing Core (NIH Grant P20 GM103638) for assistance in genome isolation, library preparation and sequencing. This work was supported by a K-INBRE postdoctoral grant to TH (NIH Grant P20 GM103418). This work was also funded by NIH Grants R00 GM114714 and R01 AI139154 to RLU.

## Data Availability

All sequencing information is available on the NCBI SRA (SRA: SRP187240), while all genomes used in this study are available on the NCBI genome database (SRA: GCA_004354385.1, GCA_005876895.1). All supplementary data used in this manuscript is available at: https://figshare.com/projects/innubila_population_genomics/87662

## Bibliography

Antunes, J. T., P. N. Leao and V. M. Vasconcelos, 2015 Cylindrospermopsis raciborskii: review of the distribution, phylogeography, and ecophysiology of a global invasive species. Frontiers in Microbiology 6: 473.

Arechederra-Romero, L., 2012 Southwest Fire Science Consortium Field Trip to the Chiricahua National Monument: Discussion of the Impacts of the 2011 Horseshoe 2 Fire, pp. in Arizona Geology Magazine, Arizona Geology Magazine.

Arizona-Geological-Survey, 2005 Arizona Geology. Arizona Geology 35: 1–6.

Astanei, I., E. Gosling, J. Wilson and E. Powell, 2005 Genetic variability and phylogeography of the invasive zebra mussel, Dreissena polymorpha (Pallas). Mol Ecol 14: 1655–1666.

Avgar, T., G. Street, and J. M. Fryxell, 2014 On the adaptive benefits of mammal migration. Canadian Journal of Zoology 92: 481–490.

Behrman, E. L., S. S. Watson, K. R. O’Brien, S. M. Heschel and P. S. Schmidt, 2015 Seasonal variation in life history traits in two Drosophila species. Journal of Evolutionary Biology 28: 1691–1704.

Buffalo, V., 2018 Scythe.

Cai, J. J., J. M. Macpherson, G. Sella and D. a. Petrov, 2009 Pervasive hitchhiking at coding and regulatory sites in humans. PLoS genetics 5: 1–13.

Chakraborty, M., R. Zhao, X. Zhang, S. Kalsow and J. J. Emerson, 2017 Extensive hidden genetic variation shapes the structure of functional elements in Drosophila. Doi.Org 50: 114967.

Chapman, J. R., T. Hill and R. L. Unckless, 2019 Balancing selection drives maintenance of genetic variation in Drosophila antimicrobial peptides. Genome Biology and Evolution 11: 2691–2701.

Charlesworth, B., D. Charlesworth and N. H. Barton, 2003 The Effects of Genetic and Geographic Structure on Neutral Variation. Annual Review of Ecology, Evolution, and Systematics 34: 99–125.

Charlesworth, B., M. T. Morgan and D. Charlesworth, 1993 The effect of deleterious mutations on neutral molecular variation. Genetics 134: 1289–1303.

Chen, X., O. Schulz-Trieglaff, R. Shaw, B. Barnes, F. Schlesinger et al., 2016 Manta: Rapid detection of structural variants and indels for germline and cancer sequencing applications. Bioinformatics 32: 1220–1222.

Cingolani, P., A. Platts, L. L. Wang, M. Coon, T. Nguyen et al., 2012 A program for annotating and predicting the effects of single nucleotide polymorphisms, SnpEff: SNPs in the genome of Drosophila melanogaster strain w1118; iso-2; iso-3. Fly 6: 80–92.

Cini, A., C. Ioriatti and G. Anfora, 2012 A review of the invasion of Drosophila suzukii in Europe and a draft research agenda for integrated pest management. Bulletin of Insectology 65: 149–160.

Cloudsley-Thompson, J. L., 1978 Human Activities and Desert Expansion. The Geographical Journal 144: 416–423.

Coe, S. J., D. M. Finch and M. M. Friggens, 2012 An Assessment of Climate Change and the Vulnerability of Wildlife in the Sky Islands of the Southwest, pp. 1–208. United States Department of Agriculture.

Crimmins, T. M., M. A. Crimmins and C. D. Bertelsen, 2011 Onset of summer flowering in a ‘Sky Island’ is driven by monsoon moisture. New Phytol 191: 468–479.

Cruickshank, T. E., and M. W. Hahn, 2014 Reanalysis suggests that genomic islands of speciation are due to reduced diversity, not reduced gene flow. Mol Ecol 23: 3133–3157.

Cutter, A. D., and B. A. Payseur, 2013 Genomic signatures of selection at linked sites: unifying the disparity among species. Nat Rev Genet 14: 262–274.

Danecek, P., A. Auton, G. Abecasis, C. A. Albers, E. Banks et al., 2011 The variant call format and VCFtools. Bioinformatics 27: 2156–2158.

DePristo, M. A., E. Banks, R. Poplin, K. V. Garimella, J. R. Maguire et al., 2011 A framework for variation discovery and genotyping using next-generation DNA sequencing data. Nature genetics 43: 491–498.

Dobzhansky, T., and B. Spassky, 1968 The Genetics of Natural Populations XL: Heterotic and Deleterious Effects of Recessive Lethals in Populations of Drosophila pseudoobscura. Genetics 59: 411–425.

Dobzhansky, T., and A. H. Sturtevant, 1937 Inversions In Chromosomes of Drosophila pseudoobscura. Genetics 23: 28–64.

Dobzhansky, T. H., A. S. Hunter, O. Pavlovsky, B. Spassky and B. Wallace, 1963 Genetics of natural populations. XXXI. Genetics of an isolated marginal population of Drosophila pseudoobscura. Genetics 48: 91–103.

Dostert, C., E. Jouanguy, P. Irving, L. Troxler, D. Galiana-Arnoux et al., 2005 The Jak-STAT signaling pathway is required but not sufficient for the antiviral response of Drosophila. Nat Immunol 6: 946–953.

Dyer, K. A., 2004 Evolutionarily Stable Infection by a Male-Killing Endosymbiont in Drosophila innubila: Molecular Evidence From the Host and Parasite Genomes. Genetics 168: 1443–1455.

Dyer, K. a., and J. Jaenike, 2005 Evolutionary dynamics of a spatially structured host-parasite association: Drosophila innubila and male-killing Wolbachia. Evolution; international journal of organic evolution 59: 1518–1528.

Dyer, K. A., M. S. Minhas and J. Jaenike, 2005 Expression and modulation of embryonic male-killing in Drosophila innubila: opportunities for multilevel selection. Evolution; international journal of organic evolution 59: 838–848.

Eilertson, K. E., J. G. Booth and C. D. Bustamante, 2012 SnIPRE: Selection Inference Using a Poisson Random Effects Model. PLoS Computational Biology 8.

Excoffier, L., M. Foll and R. J. Petit, 2009 Genetic Consequences of Range Expansions. Annual Review of Ecology, Evolution, and Systematics 40: 481–501.

Falush, D., M. Stephens and J. K. Pritchard, 2003 Inference of population structure using multilocus genotype data: Linked loci and correlated allele frequencies. Genetics 164: 1567–1587.

Fave, M. J., R. A. Johnson, S. Cover, S. Handschuh, B. D. Metscher et al., 2015 Past climate change on Sky Islands drives novelty in a core developmental gene network and its phenotype. BMC Evol Biol 15: 183.

Frichot, E., F. Mathieu, T. Trouillon, G. Bouchard and O. François, 2014 Fast and efficient estimation of individual ancestry coefficients. Genetics 196: 973–983.

Fuller, Z. L., G. D. Haynes, S. Richards and S. W. Schaeffer, 2016 Genomics of Natural Populations: How Differentially Expressed Genes Shape the Evolution of Chromosomal Inversions in. Genetics.

Gao, Z., D. Waggoner, M. Stephens, C. Ober and M. Przeworski, 2015 An estimate of the average number of recessive lethal mutations carried by humans. Genetics 199: 1243–1254.

Gillespie, J., 2004 Population Genetics: A Concise Guide. 232.

Gramates, L. S., S. J. Marygold, G. Dos Santos, J. M. Urbano, G. Antonazzo et al., 2017 FlyBase at 25: Looking to the future. Nucleic Acids Research 45: D663–D671.

Guindon, S., J.-F. Dufayard, V. Lefort, M. Anisimova, W. Hordijk et al., 2010 New algorithms and methods to estimate maximum-likelihood phylogenies: assessing the performance of PhyML 3.0. Systematic biology 59: 307–321.

Gutenkunst, R. N., R. D. Hernandez, S. H. Williamson and C. D. Bustamante, 2010 Diffusion Approximations for Demographic Inference: ∂a∂i. Nature precedings.

Haller, B. C., and P. W. Messer, 2017 asymptoticMK: A Web-Based Tool for the Asymptotic McDonald–Kreitman Test. G3: Genes, Genomes, Genetics 7: 1569–1575.

Hermisson, J., and P. S. Pennings, 2005 Soft sweeps: molecular population genetics of adaptation from standing genetic variation. Genetics 169: 2335–2352.

Hewitt, G., 2000 The genetic legacy of the Quaternary ice ages. Nature 405: 907–913.

Hill, T., B. Koseva and R. L. Unckless, 2019 The genome of Drosophila innubila reveals lineage-specific patterns of selection in immune genes. Molecular Biology and Evolution: 1–36.

Hill, T., and R. Unckless, 2020 Recurrent evolution of two competing haplotypes in an insect DNA virus. Biorxiv: 1–45.

Hoban, S., J. L. Kelley, K. E. Lotterhos, M. F. Antolin, G. Bradburd et al., 2016 Finding the Genomic Basis of Local Adaptation: Pitfalls, Practical Solutions, and Future Directions. Am Nat 188: 379–397.

Hoffmann, J. A., 2003 The immune response of Drosophila. Nature 426: 33–38.

Holmgren, K., J. A. Lee-Thorp, G. R. J. Cooper, K. Lundblad, T. C. Partridge et al., 2003 Persistent millennial-scale climatic variability over the past 25,000 years in Southern Africa. Quaternary Science Reviews 22: 2311–2326.

Http://broadinstitute.github.io/picard, Picard.

Huber, C. D., M. DeGiorgio, I. Hellmann and R. Nielsen, 2016 Detecting recent selective sweeps while controlling for mutation rate and background selection. Mol Ecol 25: 142–156.

Imler, J., and I. Elftherianos, 2009 Drosophila as a model for studying antiviral defences. Insect infection and immunity (eds Rolff J., Reyolds SE): 49–68.

Jaenike, J., and K. A. Dyer, 2008 No resistance to male-killing Wolbachia after thousands of years of infection. Journal of Evolutionary Biology 21: 1570–1577.

Jaenike, J., K. A. Dyer and L. K. Reed, 2003 Within-population structure of competition and the dynamics of male-killing Wolbachia. Evolutionary Ecology Research 5: 1023–1036.

Jaenike, J., and S. J. Perlman, 2002 Ecology and Evolution of Host-Parasite Associations: Mycophagous Drosophila and Their Parasitic Nematodes. The American Naturalist 160: S23–S39.

Joshi, N., and J. Fass, 2011 Sickle: A sliding window, adaptive, quality-based trimming tool for fastQ files. 1.33.

Kapun, M., D. K. Fabian, J. Goudet and T. Flatt, 2016 Genomic Evidence for Adaptive Inversion Clines in Drosophila melanogaster. Mol Biol Evol 33: 1317–1336.

Kimbrell, D. A., and B. Beutler, 2001 The evolution and genetics of innate immunity. Nat Rev Genet 2: 256–267.

Korneliussen, T. S., A. Albrechtsen and R. Nielsen, 2014 ANGSD: Analysis of Next Generation Sequencing Data. BMC Bioinformatics 15: 356.

Lack, J. B., C. M. Cardeno, M. W. Crepeau, W. Taylor, R. B. Corbett-Detig et al., 2015 The Drosophila genome nexus: A population genomic resource of 623 Drosophila melanogaster genomes, including 197 from a single ancestral range population. Genetics 199: 1229–1241.

Li, H., and R. Durbin, 2009 Fast and accurate short read alignment with Burrows-Wheeler transform. Bioinformatics (Oxford, England) 25: 1754–1760.

Li, H., and R. Durbin, 2011 Inference of human population history from individual whole-genome sequences. Nature 475: 493–496.

Li, H., B. Handsaker, A. Wysoker, T. Fennell, J. Ruan et al., 2009 The sequence alignment/map format and SAMtools. Bioinformatics (Oxford, England) 25: 2078–2079.

Liu, X., and Y.-X. Fu, 2015 Exploring population size changes using SNP frequency spectra. Nature genetics 47: 555–559.

Ma, X., J. L. Kelley, K. Eilertson, S. Musharoff, J. D. Degenhardt et al., 2013 Population Genomic Analysis Reveals a Rich Speciation and Demographic History of Orangutans (Pongo pygmaeus and Pongo abelii). PLoS ONE 8.

Machado, C. a., T. S. Haselkorn and M. a. F. Noor, 2007 Evaluation of the genomic extent of effects of fixed inversion differences on intraspecific variation and interspecific gene flow in Drosophila pseudoobscura and Drosophila persimilis. Genetics 175: 1289–1306.

Machado, H. E., A. O. Bergland, K. R. O’Brien, E. L. Behrman, P. S. Schmidt et al., 2015 Comparative population genomics of latitudinal variation in D. simulans and D. melanogaster. Molecular Ecology: n/a-n/a.

Mackay, T. F. C., S. Richards, E. a. Stone, A. Barbadilla, J. F. Ayroles et al., 2012 The Drosophila melanogaster genetic reference panel. Nature 482: 173–178.

Manthey, J. D., and R. G. Moyle, 2015 Isolation by environment in White-breasted Nuthatches (Sitta carolinensis) of the Madrean Archipelago sky islands: a landscape genomics approach. Mol Ecol 24: 3628–3638.

Marinkovic, D., 1967 Genetic Loads Affecting Fecundity in Natural Populations of Drosophila pseudoobscura. Genetics: 61-71.

Markow, T. A., and P. O’Grady, 2006 Drosophila: a guide to species identification.

Martin, M., 2011 Cutadapt removes adapter sequences from high-throughput sequencing reads. Technical Notes: 1–12.

Marzo, M., M. Puig and A. Ruiz, 2008 The Foldback-like element Galileo belongs to the P superfamily of DNA transposons and is widespread within the Drosophila genus. Proceedings of the National Academy of Sciences of the United States of America 105: 2957–2962.

Matthey-Doret, R., and M. C. Whitlock, 2018 Background selection and the statistics of population differentiation: consequences for detecting local adaptation. Biorxiv: 1–5.

McCormack, J. E., H. Huang and L. L. Knowles, 2009 Sky Islands, pp. 841-843 in Encyclopedia of islands.

McDonald, J. H., and M. Kreitman, 1991a Adaptive protein evolution at the Adh locus in Drosophila. Nature 351: 652–654.

McDonald, J. H., and M. Kreitman, 1991b Adaptive protein evolution at the Adh locus in Drosophila. Nature 351: 652–654.

McKenna, A., M. Hanna, E. Banks, A. Sivachenko, K. Cibulskis et al., 2010 The Genome Analysis Toolkit: A MapReduce framework for analyzing next-generation DNA sequencing data. Proceedings of the International Conference on Intellectual Capital, Knowledge Management & Organizational Learning 20: 1297–1303.

McVean, G., 2007 The structure of linkage disequilibrium around a selective sweep. Genetics 175: 1395–1406.

Merkling, S. H., and R. P. van Rij, 2013 Beyond RNAi: Antiviral defense strategies in Drosophila and mosquito. Journal of Insect Physiology 59: 159–170.

Messer, P. W., and D. A. Petrov, 2012 The McDonald-Kreitman Test and its Extensions under Frequent Adaptation: Problems and Solutions. Proceedings of the National Academy of Sciences 110: 8615–8620.

Messer, P. W., and D. A. Petrov, 2013 Population genomics of rapid adaptation by soft selective sweeps. Trends in Ecology & Evolution 28: 659–669.

Misztal, L. W., G. Garfin and L. Hansen, 2013 Responding to Climate Change Impacts in the Sky Island Region: From Planning to Action. USDA Forest Service Proceedings P67.

Narasimhan, V., P. Danecek, A. Scally, Y. Xue, C. Tyler-Smith et al., 2016 BCFtools/RoH: A hidden Markov model approach for detecting autozygosity from next-generation sequencing data. Bioinformatics 32: 1749–1751.

Nei, M., 1987 Molecular evolutionary genetics. Columbia university press.

Nei, M., and J. Miller, 1990 A Simple Method for Estimating Average Number of Nucleotide Substitutions Within and Between Populations From Restriction Data. Genetics 125: 873–879.

Nielsen, R., 2005 Molecular signatures of natural selection. Annu Rev Genet 39: 197–218.

Noor, M. a. F., D. a. Garfield, S. W. Schaeffer and C. a. Machado, 2007 Divergence between the Drosophila pseudoobscura and D. persimilis genome sequences in relation to chromosomal inversions. Genetics 177: 1417–1428.

Oneal, E., D. B. Lowry, K. M. Wright, Z. Zhu and J. H. Willis, 2014 Divergent population structure and climate associations of a chromosomal inversion polymorphism across the Mimulus guttatus species complex. Mol Ecol 23: 2844–2860.

Palmer, W. H., J. Joosten, G. J. Overheul, P. W. Jansen, M. Vermeulen et al., 2018 Induction and suppression of NF-κB signalling by a DNA virus of Drosophila.

Parmesan, C., and G. Yohe, 2003 A globally coherent fingerprint of climate change impacts across natural systems. Nature 421: 37–42.

Patterson, J. T., 1954 Studies in the genetics of Drosophila., pp.

Patterson, J. T., and W. S. Stone, 1949 Studies in the genetics of Drosophila., pp. 7–17 in University of Texas Publications.

Perlman, S. J., G. S. Spicer, D. Dewayne Shoemaker and J. Jaenike, 2003 Associations between mycophagous Drosophila and their Howardula nematode parasites: A worldwide phylogenetic shuffle. Molecular Ecology 12: 237–249.

Pool, J., and C. H. Langley, 2013 DPGP3.

Pool, J. E., R. B. Corbett-detig, R. P. Sugino, K. A. Stevens, C. M. Cardeno et al., 2012 Population Genomics of Sub-Saharan Drosophila melanogaster: African Diversity and Non-African Admixture. PLoS Genetics 8: 1–24.

Porretta, D., V. Mastrantonio, R. Bellini, P. Somboon and S. Urbanelli, 2012 Glacial history of a modern invader: phylogeography and species distribution modelling of the Asian tiger mosquito Aedes albopictus. PLoS One 7: e44515.

Przeworski, M., G. Coop, J. D. Wall and S. Url, 2011 The Signature of Positive Selection on Standing Genetic Variation. Evolution 59: 2312–2323.

Puzey, J. R., J. H. Willis and J. K. Kelly, 2017 Population structure and local selection yield high genomic variation in Mimulus guttatus. Mol Ecol 26: 519–535.

Rankin, M. A., and J. C. A. Burchsted, 1992 The Cost of Migration in Insects. Annual Review of Entymology 37: 533–559.

Rastogi, S., and D. a. Liberles, 2005 Subfunctionalization of duplicated genes as a transition state to neofunctionalization. BMC evolutionary biology 5: 28.

Rausch, T., T. Zichner, A. Schlattl, A. M. Stutz, V. Benes et al., 2012 DELLY: structural variant discovery by integrated paired-end and split-read analysis. Bioinformatics 28: i333–i339.

Rodriguez-Trelles, F., G. Alvarez and C. Zapata, 1996 Time-Series Analysis of Seasonal Changes of the 0 Inversion polymorphism of Drosqphiila subobscura. Genetics 142: 179–187.

Rosenzweig, C., D. Karoly, M. Vicarelli, P. Neofotis, Q. Wu et al., 2008 Attributing physical and biological impacts to anthropogenic climate change. Nature 453: 353–357.

Schrider, D. R., D. Houle, M. Lynch and M. W. Hahn, 2013 Rates and genomic consequences of spontaneous mutational events in Drosophila melanogaster. Genetics 194: 937–954.

Scott Chialvo, C. H., and T. Werner, 2018 Drosophila, destroying angels, and deathcaps! Oh my! A review of mycotoxin tolerance in the genus Drosophila. Frontiers in Biology 13: 91–102.

Searle, J. B., P. Kotlik, R. V. Rambau, S. Markova, J. S. Herman et al., 2009 The Celtic fringe of Britain: insights from small mammal phylogeography. Proc Biol Sci 276: 4287–4294.

Sella, G., D. A. Petrov, M. Przeworski and P. Andolfatto, 2009 Pervasive natural selection in the Drosophila genome? PLoS Genet 5: e1000495.

Shoemaker, D. D., V. Katju and J. Jaenike, 1999 Wolbachia and the evolution of reproductive isolation between Drosophila recens and Drosophila subquinaria. Evolution 53: 1157–1164.

Smit, A. F. A., and R. Hubley, 2013-2015 RepeatMasker Open-4.0, pp. RepeatMasker.

Smith, C. I., and B. D. Farrell, 2005 Phylogeography of the longhorn cactus beetle Moneilema appressum LeConte (Coleoptera: Cerambycidae): was the differentiation of the Madrean sky islands driven by Pleistocene climate changes? Mol Ecol 14: 3049–3065.

Smith, F. A., J. L. Betancourt and J. H. Brown, 1995 Evolution of Body Size in the Woodrat Over the Past 25,000 Years of Climate Change. Science 270: 2012–2014.

Smith, N. G. C., and A. Eyre-Walker, 2002 Adaptive protein evolution in Drosophila. Nature 415: 1022–1024.

Stajich, J. E., and M. W. Hahn, 2005 Disentangling the effects of demography and selection in human history. Mol Biol Evol 22: 63–73.

Stoletzki, N., and A. Eyre-Walker, 2011 Estimation of the neutrality index. Molecular Biology and Evolution 28: 63–70.

Subramanian, A., P. Tamayo, V. K. Mootha, S. Mukherjee, B. L. Ebert et al., 2005 Gene set enrichment analysis: A knowledge-based approach for interpreting genome-wide expression profiles. PNAS 102: 15545–15550.

Tajima, F., 1989 Statistical method for testing the neutral mutation hypothesis by DNA polymorphism. Genetics 123: 585–595.

Takeda, K., and S. Akira, 2005 Toll-like receptors in innate immunity. International Immunology 17: 1–14.

Tigano, A., and V. L. Friesen, 2016 Genomics of local adaptation with gene flow. Mol Ecol 25: 2144–2164.

Unckless, R. L., 2011 A DNA Virus of Drosophila. PLoS ONE 6: e26564.

Unckless, R. L., and J. Jaenike, 2011 Maintenance of a Male-Killing Wolbachia in Drosophila innubila By Male-Killing Dependent and Male-Killing Independent Mechanisms. Evolution 66: 678–689.

Vicoso, B., and D. Bachtrog, 2015 Numerous Transitions of Sex Chromosomes in Diptera. PLoS Biology 13: 1–22.

Walsh, D. B., M. P. Bolda, R. E. Goodhue, A. J. Dreves, J. Lee et al., 2011 Drosophila suzukii (Diptera: Drosophilidae): Invasive Pest of Ripening Soft Fruit Expanding its Geographic Range and Damage Potential. Journal of Integrated Pest Management 2: 1–7.

Watanabe, T. K., O. Yamaguchi and T. Mukai, 1974 The Genetic Variability of Third Chromosomes in a Local Population of Drosophila melanogaster. Genetics 82: 63–82.

Weir, B. S., and C. C. Cockerham, 1984 Estimating F-Statistics for the Analysis of Population Structure. Evolution 38: 1358–1370.

Werren, J. H., L. Baldo and M. E. Clark, 2008 Wolbachia: master manipulators of invertebrate biology. Nat Rev Microbiol 6: 741–751.

White, T. A., S. E. Perkins, G. Heckel and J. B. Searle, 2013 Adaptive evolution during an ongoing range expansion: the invasive bank vole (Myodes glareolus) in Ireland. Mol Ecol 22: 2971–2985.

Wiens, J. J., A. Camacho, A. Goldberg, T. Jezkova, M. E. Kaplan et al., 2019 Climate change, extinction, and Sky Island biogeography in a montane lizard. Mol Ecol 28: 2610–2624.

Wright, S. I., B. Lauga and D. Charlesworth, 2003 Subdivision and haplotype structure in natural populations of Arabidopsis lyrata. Molecular Ecology 12: 1247–1263.

Ye, K., M. H. Schulz, Q. Long, R. Apweiler and Z. Ning, 2009 Pindel : a pattern growth approach to detect break points of large deletions and medium sized insertions from paired-end short reads. Bioinformatics 25: 2865–2871.

Zambon, R. A., M. Nandakumar, V. N. Vakharia and L. P. Wu, 2005 The Toll pathway is important for an antiviral response in Drosophila. Proceedings of the National Academy of Sciences 102: 7257–7262.

