## Supplementary Data for "Adaptation, ancestral variation and gene flow in a ‘Sky Island’ *Drosophila* species"

### Supplementary Methods

We used dnaPipeTE (GOUBERT *et al.* 2015) to quantify the extent that repetitive element content differed across the populations. Our approach assumed a genome size of 168Mbp, with the number of randomly sampled reads equal to 1-fold coverage of the genome, resampling each strain 2 times to get the average estimate of each strains TE content. Following TE identification, we grouped sequences by known super-families and compared the proportion of the genome composed of each superfamily across strains in the populations. We also used a reciprocal blast (e-value < 0.00000001) (ALTSCHUL *et al.* 1990) to identify TE families present in each strain.

We confirmed TE families shared between the previous RepeatModeler (SMIT AND HUBLEY 2008) annotation of the *D. innubila* reference genome and the dnaPipeTE annotation using blast (e-value < 10e-08) (ALTSCHUL *et al.* 1990). After confirming that they did not differ in content, we called TE insertions in each strain across the genome using PopoolationTE2 (KOFLER *et al.* 2016), then merged the output and calculated the frequency of insertions, grouping by TE order, population and if the insertion was exonic, intronic, non-coding or flanking a gene (500bp up or downstream of start or end). When considering individual TE families in *D. innubila*, we used the RepBase TE names and identifications (BAO *et al.* 2015).

### Supplementary Results

*Evidence for divergence in the X chromosome over time*

We next compared the samples from the 35 CH males in 2017 to those we sequenced from a CH collection of 38 males in September 2001 to identify changes over time between populations (due to elevated $F_{ST}$ because of differences in allele frequencies between populations). We find little differentiation between the two timepoints (median $F_{ST}$ = 0.0004, 99$^{th}$ percentile = 0.0143), and find no significant enrichments (GO *p*-value < 0.05) in the upper 97.5$^{th}$ percentile of $F_{ST}$. However, $F_{ST}$ is higher (but still low) in the genes at the telomere of the X chromosome (Supplementary Figure 10A, Muller element A, 35-40.5Mb, median $F_{ST}$ = 0.0029). As X chromosome coverage is significantly higher in 2001 samples compared to 2017 samples (Wilcoxon Rank Sum Text W = 385, *p*-value = 0.002461), we reasoned that we are more likely to call SNPs in 2001 than 2017 and so could create false signatures of differentiation with this. Because of that, we down sampled all samples to equal coverage and again called variation again and estimated $F_{ST}$. Again, we find $F_{ST}$ is higher in the X telomere, suggesting a true difference in allele frequencies between timepoints (Supplementary Figure 10A). Looking at actual allele frequency differences between time points using synonymous variation, the minor allele frequency increases between 2001 and 2017 at the X telomere while most other chromosomes appear to show little change (Supplementary Figure 10B). Interestingly, there is also evidence of recent selective sweeps in the telomere of X (Supplementary Figure 9A). The minor allele frequency also decreases on average on Muller element B between 2001 and 2017 (Supplementary Figure

10B). This suggests something else may be influencing allele frequency change on Muller B compared to other autosomes.

We see little evidence of divergence between samples collected in 2001 and 2017 (Supplementary Figure 10A), though exceptions are found on Muller B and the telomeric end of the X chromosome (Supplementary Figure 10B). Selection is likely polygenic and relatively weak genome wide, resulting in very few changes in allele frequency in the 16 years (~80 generations) between samples. Both divergent regions are associated with reduced recombination, where hitchhiking can occur, and differentiation can be more pronounced, allowing us to identify differences between timepoints even with weak selection (HERMISSON AND PENNINGS 2005; MCVEAN 2007; PRZEWORSKI *et al.* 2011). Segregating inversions, such as those on Muller element B, are frequently associated with local or temporal variation (DOBZHANSKY AND STURTEVANT 1937; RODRIGUEZ-TRELLES *et al.* 1996; ONEAL *et al.* 2014), additionally there was an extensive forest fire in 2011 which could plausibly have been a strong selective force driving a change in allele frequency on Muller B between time points (Supplementary Figure 10B) (ARECHEDERRA-ROMERO 2012). The temporal divergence limited to the X telomere (Supplementary Figure 10A) may be associated with a X-linked selfish element driving divergence over time instead (BURT AND TRIVERS 2006), and selfish elements frequently accumulate in non-recombining regions to prevent the breakdown of synergistic genetic components (BURT AND TRIVERS 2006). Other selfish elements shown a high rate of fixation of mutations resulting in elevated divergence compared to the rest of the genome (BRAND *et al.* 2015), which could be causing the temporal divergence seen in *D. innubila* X-telomere (Supplementary Figure 10). Previous work has also shown that an X chromosome meiotic driver can lower the frequency of *Wolbachia* infection and cause *Wolbachia* extinction, specifically for a male-killing *Wolbachia* as is found in *D. innubila* (ENGELSTADTER *et al.* 2004), given this it’s possible the X-telomere could be evolving in response to the *Wolbachia* to attempt to resist the male-killing effects of the *Wolbachia* or drive it to extinction (ENGELSTADTER *et al.* 2004).

*Transposable element content of D. innubila across multiple populations*

We characterized the repetitive content across our samples using dnaPipeTE (GOUBERT *et al.* 2015) and called TE insertions per line using PopoolationTE2 (KOFLER *et al.* 2016). The reference *D. innubila* genome contains 154 different TE families along with varying satellites and simples repeats, with resequenced individuals varying from 4.4% to 38.4% of reads matching repetitive sequences. Strains varied from 1913 to 7479 TE insertions per strain in the non-repetitive portion of the genome. Like nuclear polymorphism (Supplementary Figure 1), we find little population structure by shared TE insertions, though strains do seem to disperse primarily by the number of insertions (Supplementary Figure 12B).

Similar to *D. melanogaster* (CHARLESWORTH AND LANGLEY 1989; CHARLESWORTH *et al.* 1997; PETROV *et al.* 2011; KOFLER *et al.* 2012; KOFLER *et al.* 2015), *D. innubila* harbors a significant excess of low frequency TE insertions compared to the SFS of synonymous variants (Supplementary Figure 12A, GLM Count ~ Frequency * SNP or TE, t-value = -16.401, p-value = 1.889e-60), with no difference in the insertion frequency spectra between populations (GLM Count ~ Frequency * TE order * Population, t-value = -0.341, p-value = 0.733). This implies in every population, TE insertions are on average mildly deleterious and removed via purifying selection (CHARLESWORTH AND LANGLEY 1989).

Using dnaPipeTE (GOUBERT *et al.* 2015), we find a significantly higher density of RC & TIR elements compared to other repeat orders (Supplementary Figure 12C, t-value = 3.555 p-value = 3.745e-04), consistent with the reference genome (HILL *et al.* 2019). The density of repetitive content is also higher genome wide in the CH and PR populations compared to HU and SR (Supplementary Figures 12C & D, t-value = 2.856, p-value = 4.291e-03). This is in keeping with a more recent bottleneck for these species reducing effective population size and efficacy of selection, resulting in bursts of repeat activity with relaxed selection for removal of insertions. These changes are primarily driven by an expansion of simple repeats in the CH population (Supplementary Figure 12D, GLM t-value = 3.978, p-value = 7.31e-05) and an expansion of TIR elements in the PR population (Supplementary Figure 12D, GLM t-value = 3.914, p-value = 9.52e-05). Specifically, we see expansions of the satellite CASAT_HD (GLM t-value = 5.554, p-value = 8.832e-08) and the simple repeat sequences CAACAA, CTC and GTGT in the CH population when compared to all other populations (GLM t-value = 9.204, p-value = 2.555e-17). In the PR population we find significantly higher abundances of a TE families closely related to *Tetris_Dvir* (GLM t-value = 13.641, p-value = 2.889e-32), *Helitron-2N1_DVir* (GLM t-value = 12.381, p-value = 2.789e-28) and *Chapaev3-1_PM* (GLM t-value = 11.472, p-value = 1.662e-24) compared to other populations. We do not find any evidence that particular TE orders are more abundant on any one chromosome in *D. innubila* (GLM t-value = 1.854, p-value = 0.633), though do find TEs are at significantly higher insertion densities in the inverted regions of Muller element A than at the regions of the genome (Wilcoxon Rank Sum Test W= 19763, *p*-value = 0.01488). This suggests the lack of recombination in the inverted region is allowing the accumulation of repetitive content on Muller element A.

TE insertions are usually assumed to be at least mildly deleterious (CHARLESWORTH AND LANGLEY 1989; PETROV *et al.* 2011). In *D. innubila*, TE density is lower in regions flanking genes or within genes compared to non-coding regions (GLM t-value = -6.538, p-value = 6.23e-11), consistent with the deleterious assumption. However, the frequency of TE insertions was significantly higher in exonic regions compared to introns and UTRs (Supplementary Figure 12A, GLM t-value = 4.040, p-value = 5.34e-05), across all populations, which we may have observed as these are wild caught flies and so may have more recessive deleterious insertions segregating in the population than are seen in inbred samples. Overall the repetitive content in *Drosophila innubila* appears to be mildly deleterious, with TE insertions shared between locations by migration. Despite this there are some major differences in the repeat content of each population, possibly due to the stochastic effect of population bottlenecks.

This may have occurred due to a founder effect following the population bottleneck, where a majority of CH founders by chance had a higher proportion of particular satellites or simple repeats (CHARLESWORTH *et al.* 2003). Alternatively, the bottleneck could have fixed segregating recessive variation which limits the regulation of repetitive content in the genome, leading to its expansion. However, if this was the case and satellite expansion is even mildly deleterious, we would expect migratory rescue of repeat regulation machinery. A third possibility is that satellite expansion is associated with local evolutionary dynamics either involved in adaptation or genetic conflict (GARRIDO-RAMOS 2017; LOWER *et al.* 2018).

**Supplementary Figure 1: A.** Population size history of *Drosophila innubila* backwards in time for each population. **B.** Population size history on the Log10 scale of *Drosophila innubila* backwards in time for each population. **C.** Results of Structure software (FALUSH *et al.* 2003) for estimating population structure between locations for 100,000 sampled synonymous polymorphisms from all autosomes, with a K=3 (estimated optimal K value). Note that this plot summarizes all autosomes (excluding Muller B) and the X chromosome due to very little structure between locations for all chromosomes. **D.** Results of Structure software (FALUSH *et al.* 2003) for estimating population structure between locations for 16 polymorphisms on the mitochondria, with a K=3 (estimated optimal K value).

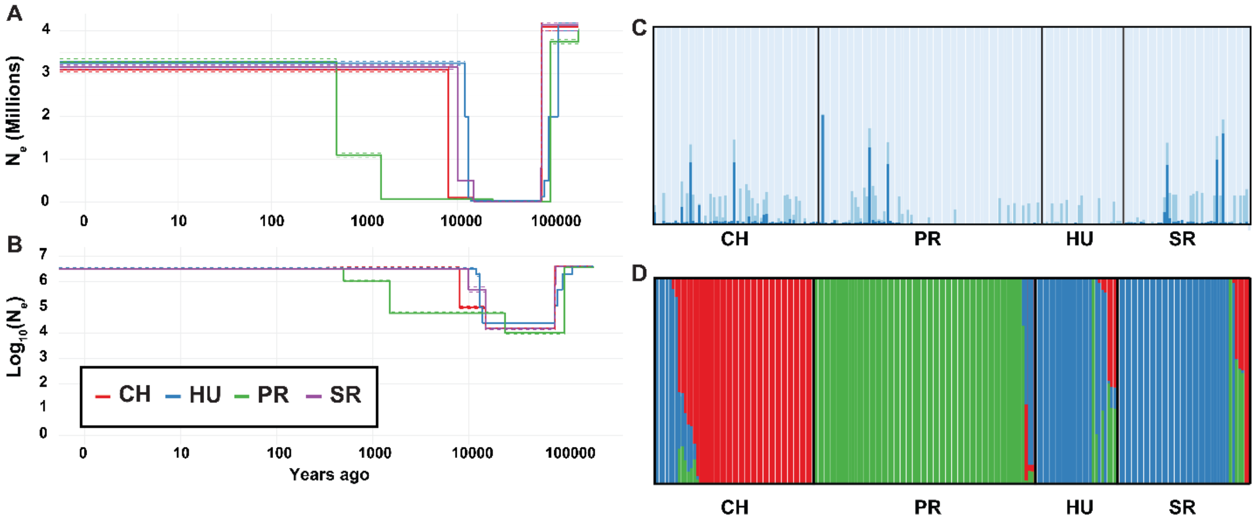

**Supplementary Figure 2:** The proportion of SNPs across the genome which are exclusive to a single population, split by populations. SNPs are shown in windows of 1kbp. The upper 97.5th percentile windows are colored red.

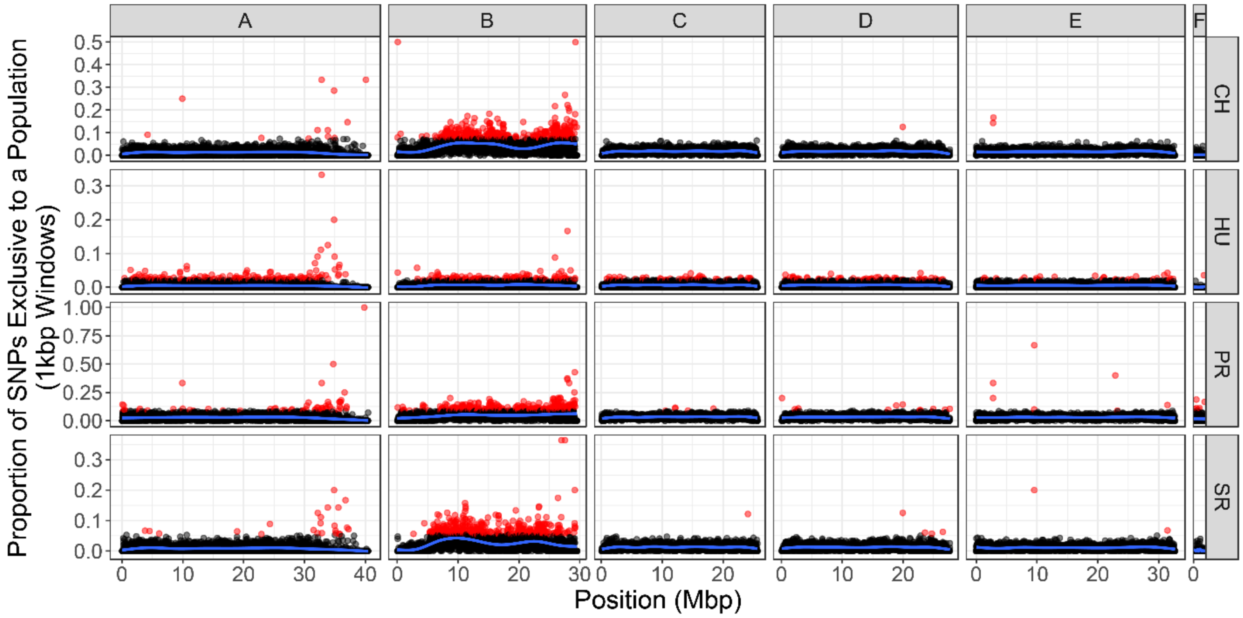


**Supplementary Figure 3:** $F_{ST}$ by gene across all Muller elements for each population, located by loci (in bp) on the Muller element.

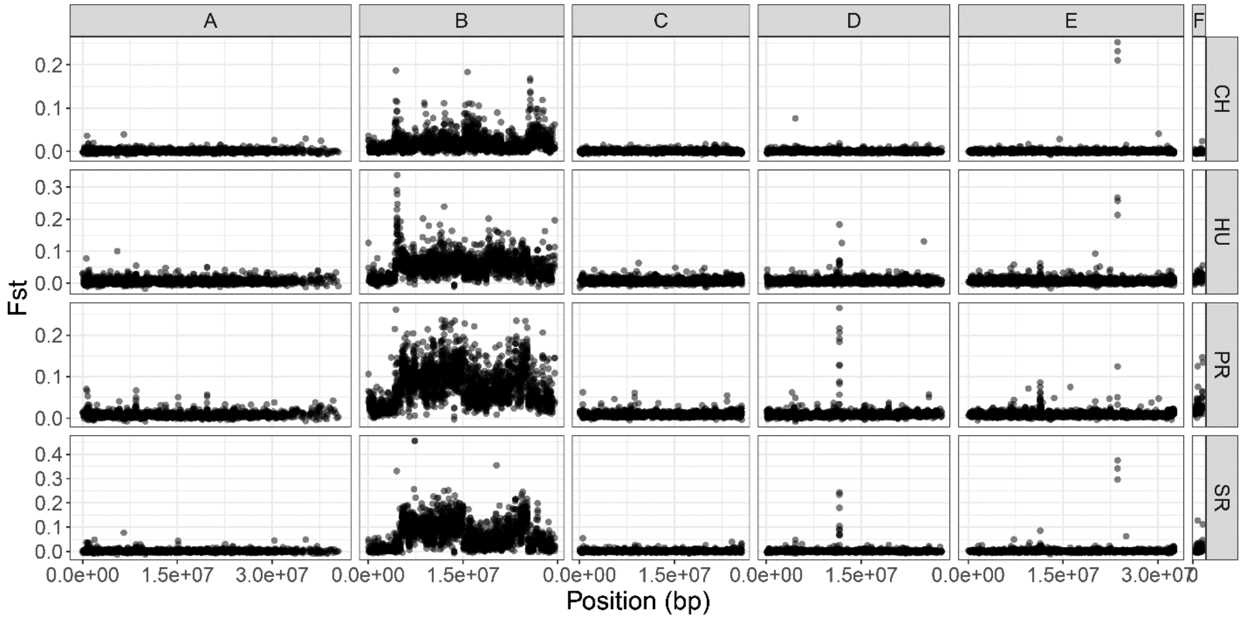

**Supplementary Figure 4: A.** Pairwise $F_{ST}$ comparison between populations, by gene across all Muller elements for each population, located by loci (in bp) on the Muller element. **B.** Pairwise $D_{XY}$ comparison between populations, by gene across all Muller elements for each population, located by loci (in bp) on the Muller element. The upper 97.5th percentile windows are colored red.

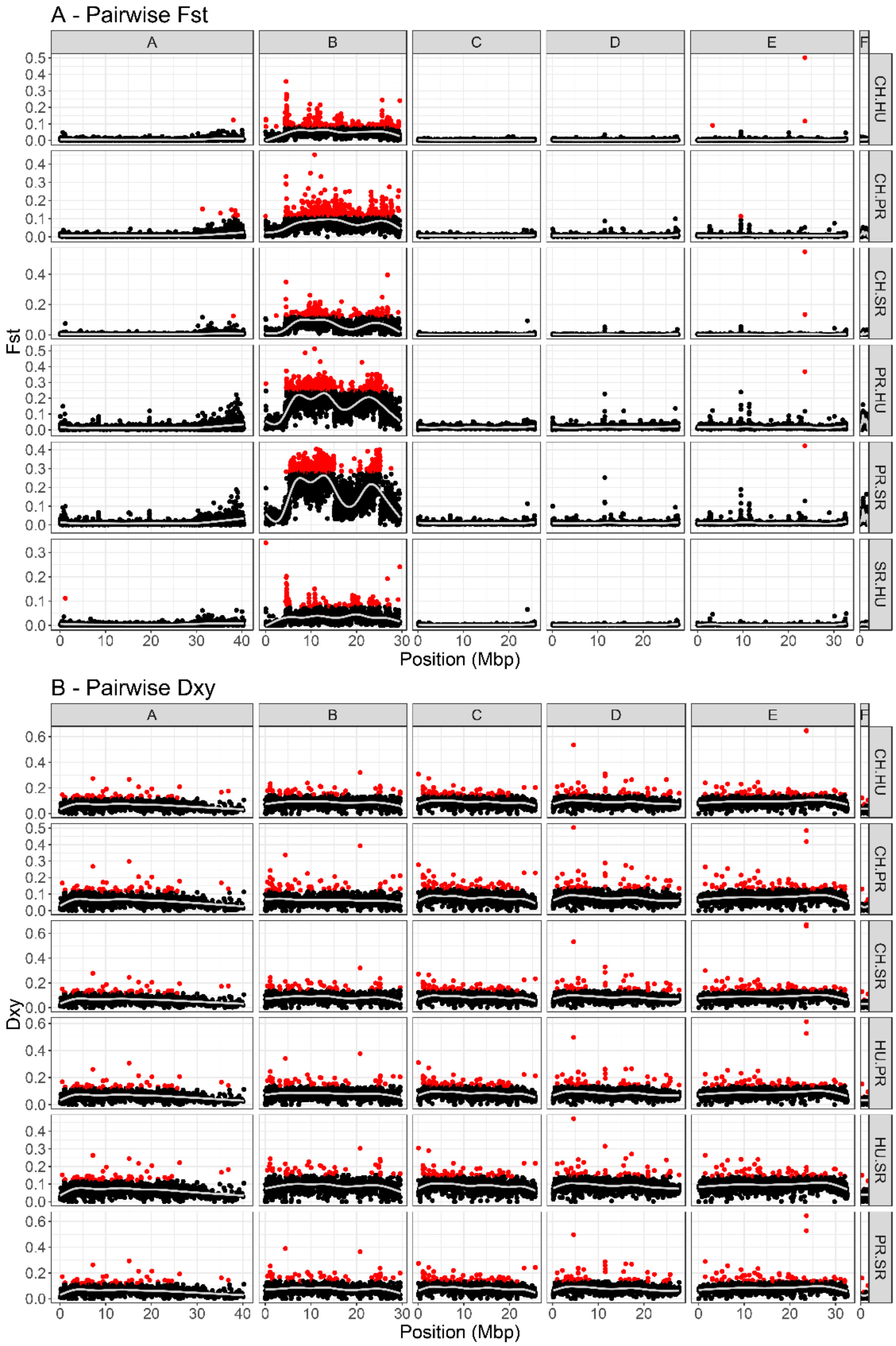

**Supplementary Figure 5:** Tajima's D by gene across all Muller elements for each population, located by loci (in bp) on the Muller element. The grey dashed line shows a Tajima's D of 0.

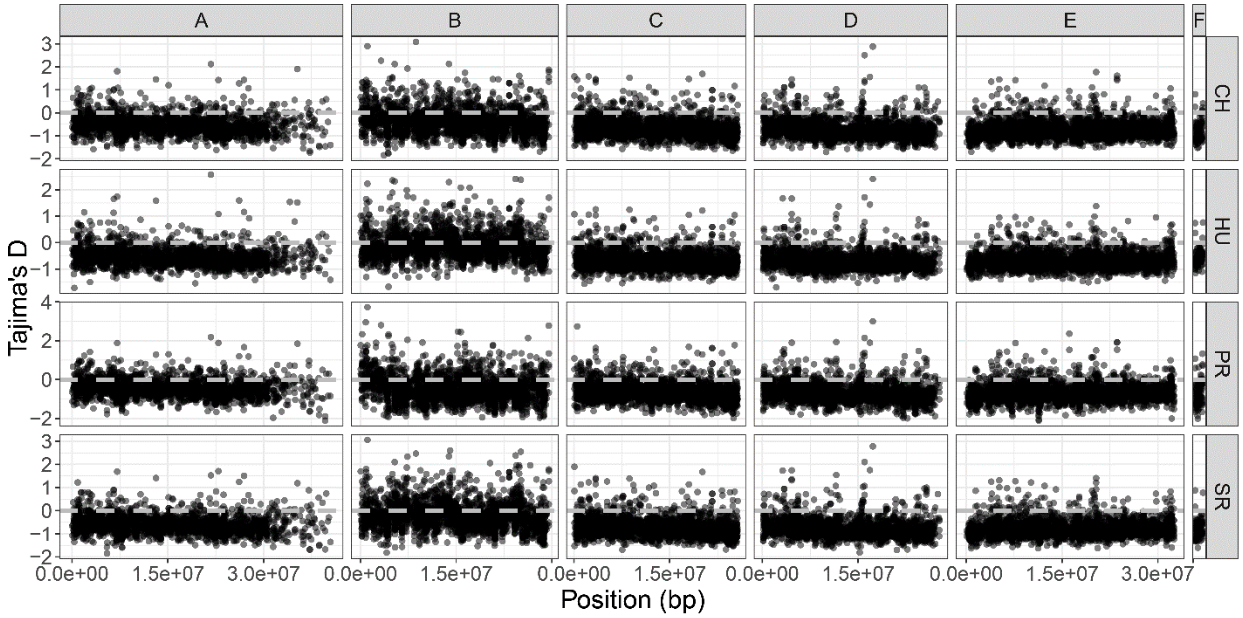

**Supplementary Figure 6:** Population genetic statistics (Pairwise diversity, Tajima's D and $F_{ST}$) for genes in each immune category, for each population. Each plot has a dotted line to show the genomic background statistics for each population. The Tajima's D plot contains a dashed line to show 0.

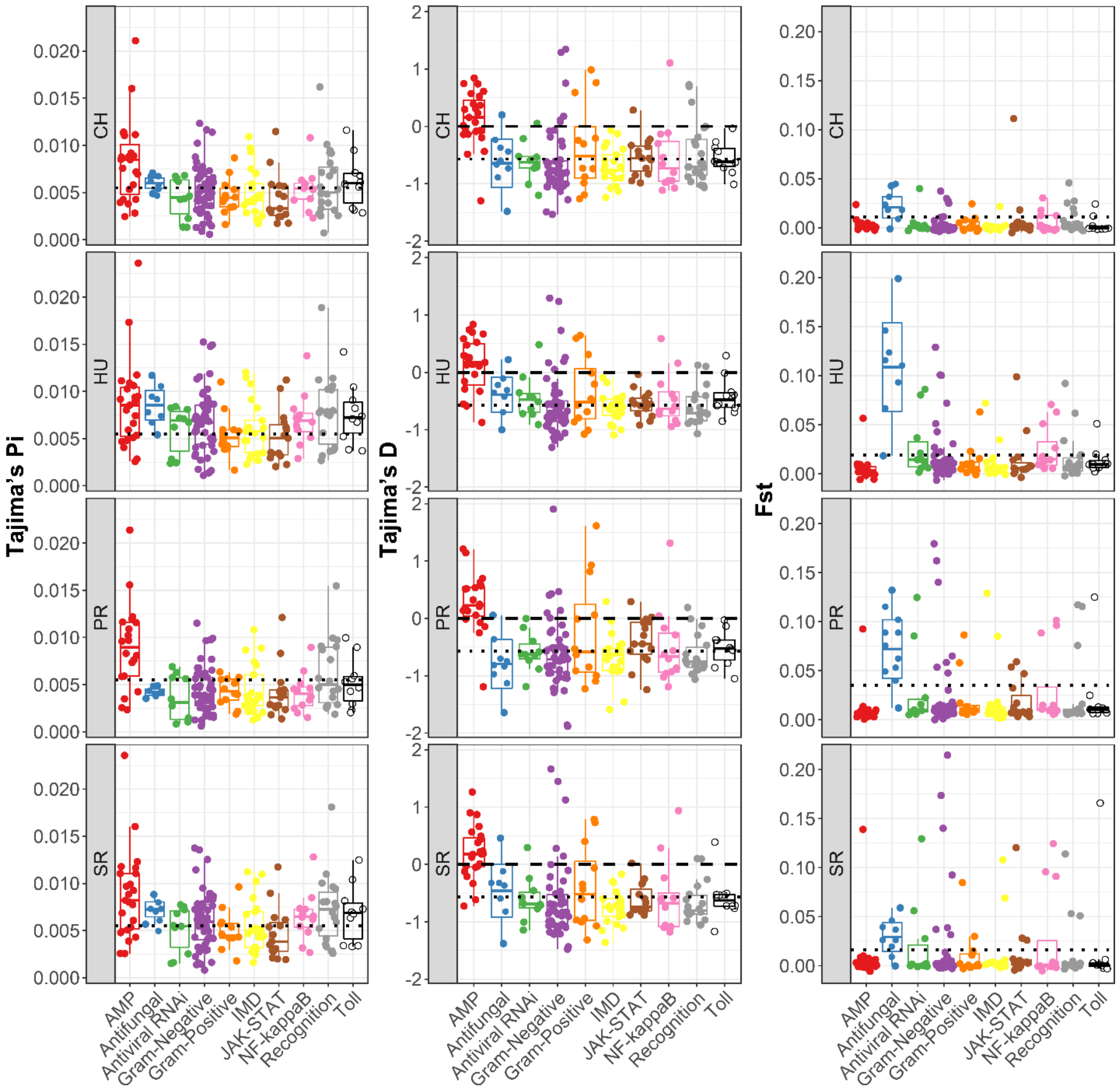

**Supplementary Figure 7:** Mean copy number per population for genes of interest in $F_{ST}$ peaks.

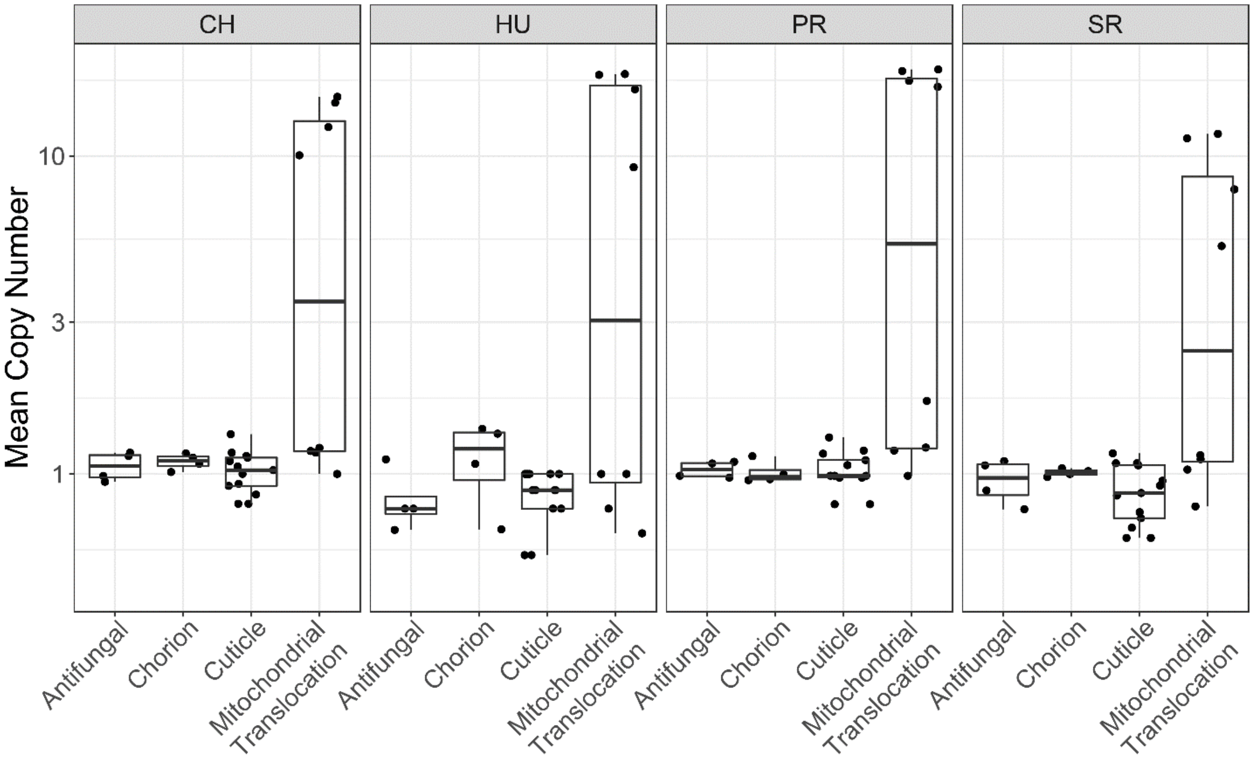

**Supplementary Figure 8: A.** Within population pairwise diversity per gene across the *D. innubila* genome. **B.** $D_{XY}$ per gene for each population, compared to all other populations. In both plots, the upper 97.5th percentile genes are marked in red, while genes below the 97.5th percentile is marked in black.

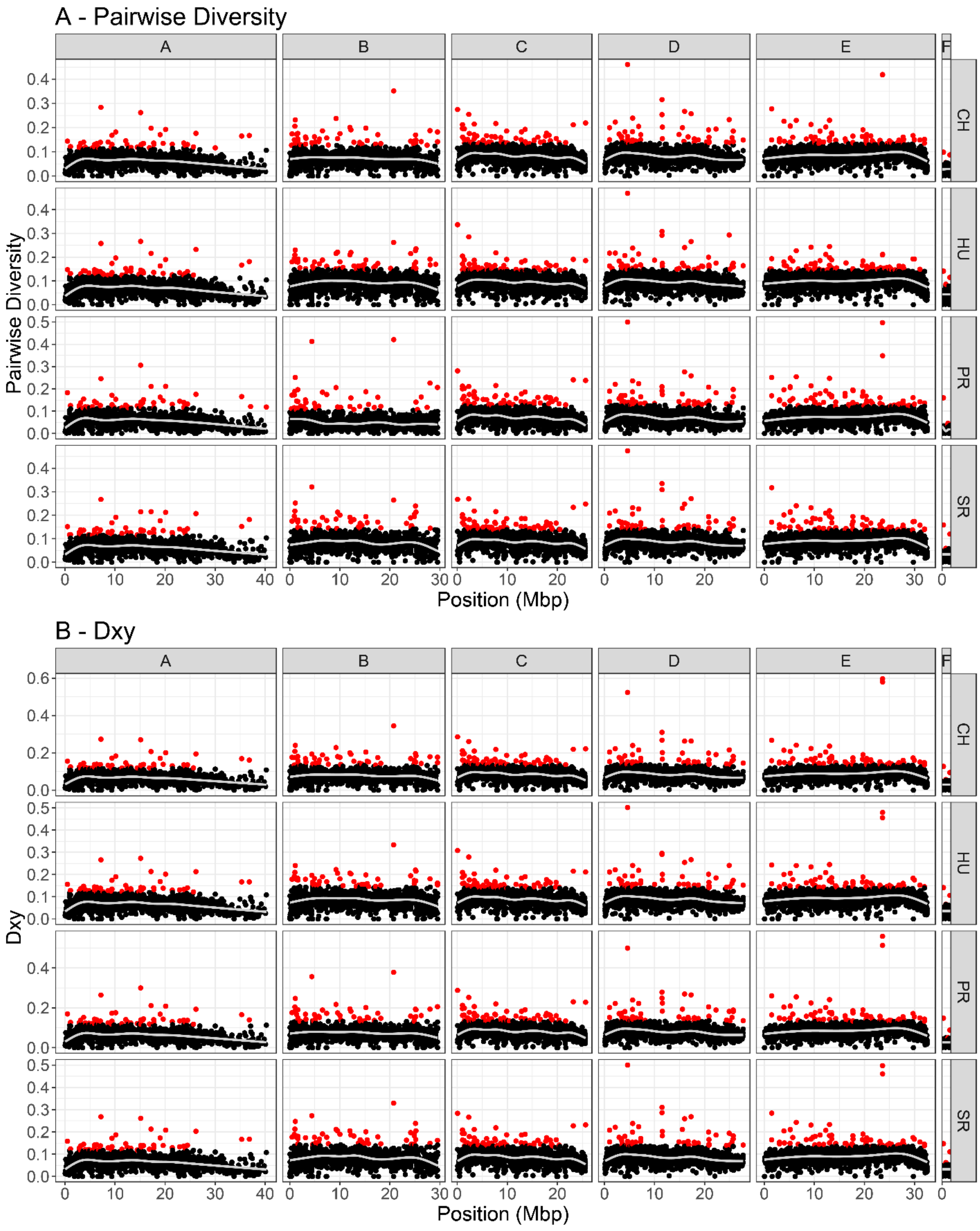

**Supplementary Figure 9:** Composite likelihood score for a selective sweep in 1kbp windows of the genome estimated using Sweepfinder2. Separated by chromosome and population. **A.** Genome wide composite likelihood score. **B.** Focus on 18.5-19Mbp of Muller element D, to show strongest selective sweep in each the PR population.

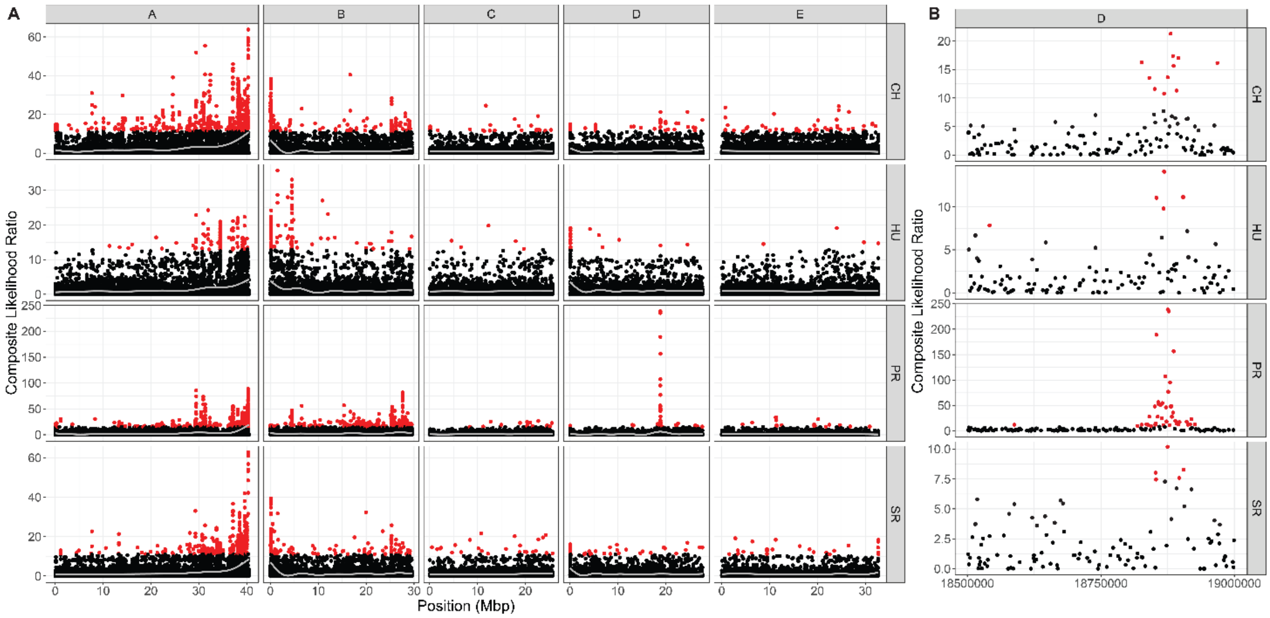

**Supplementary Figure 10: A.** $F_{ST}$ of genes between CH samples from 2001 and 2017, by chromosome and position. We also compared FST between timepoints using variation for all samples at their sequenced coverage and with all samples downsampled to the same coverage. **B.** Minor allele frequency difference curve across the genome (averaged over 2000 SNPs, sliding 1000 SNPs) between 2001 Male Chiricahua and 2017 Male Chiricahua samples. Shows average difference in the minor allele frequencies (based on total 2017 sample).

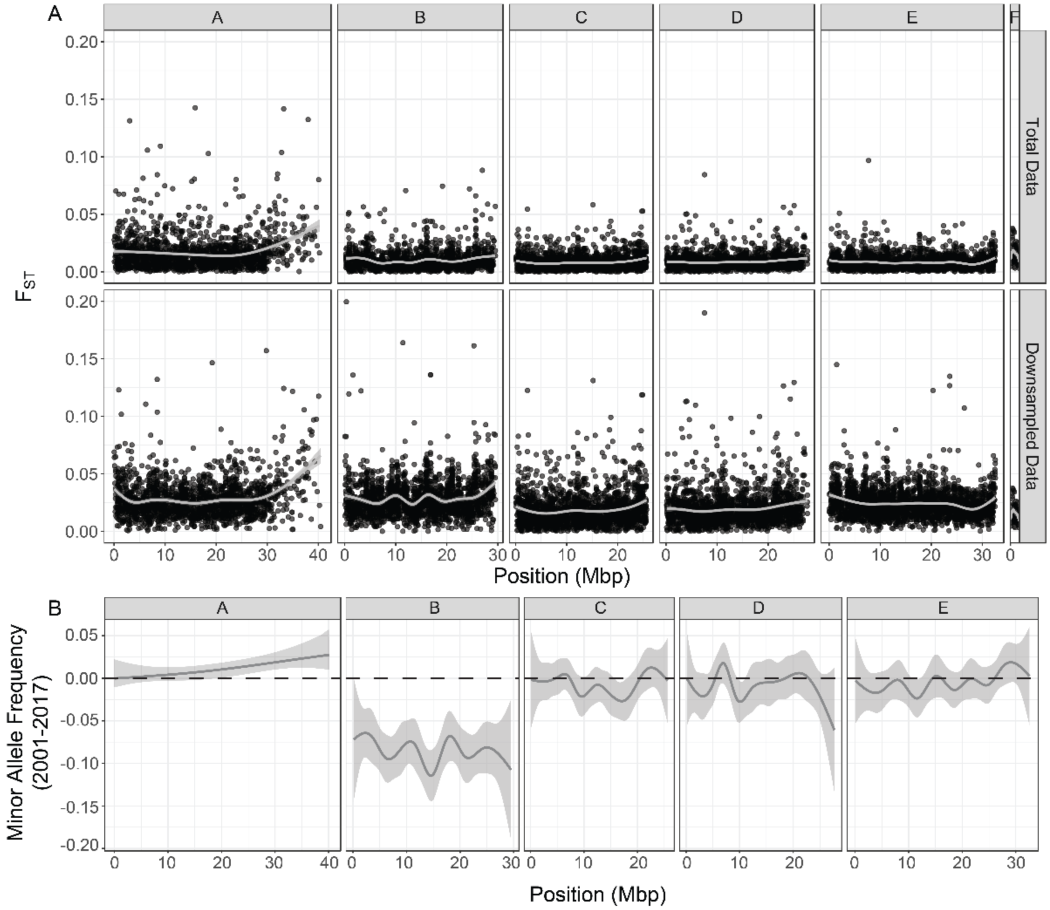

**Supplementary Figure 11:** Asymptotic α for immune categories, cuticle development proteins and all proteins, with 95% confidence intervals for categories. Categories marked with a * are significantly higher than the background following a permutation test (<0.05). 0 is marked with a dashed line. In the 'All' category, the median and 95% confidence interval is calculated across all functional categories, while in the specific functional categories the 95% confidence intervals for Asymptotic α are calculated using AsymptoticMK (MESSER AND PETROV 2012; HALLER AND MESSER 2017).

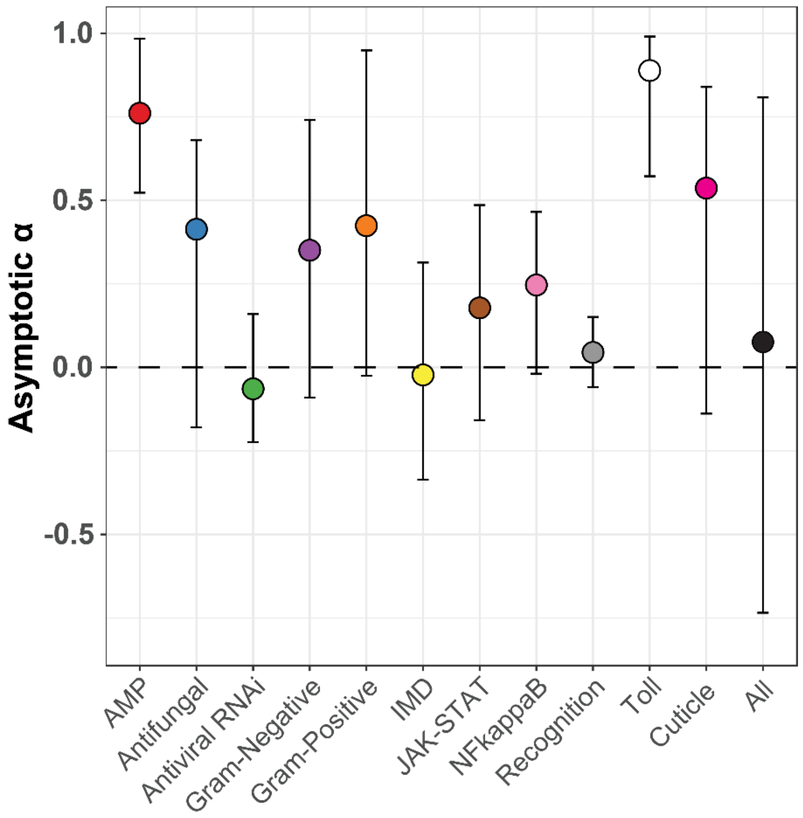


**Supplementary Figure 12:** The transposable element content of *Drosophila innubila*. **A.** Insertion frequency spectra for TEs in *D. innubila* separated by TE order and the location of insertion (e.g. coding region, non-coding, intronic). **B.** Principle component analysis (showing PCs 1 & 2) of TE insertions across *D. innubila* strains shows little population structure. Strains are labelled by their population (both shape and color). **C.** Mean TE insertion density per 1Mb window (sliding 1Mb) for each population of *D. innubila,* identified using PopoolationTE2. TE insertions are colored by their order. **D.** Proportion of the genome made up of repetitive content for each strain, as found with dnaPipeTE. Strains are ordered by total TE content from most to least, with bars colored by TE order.

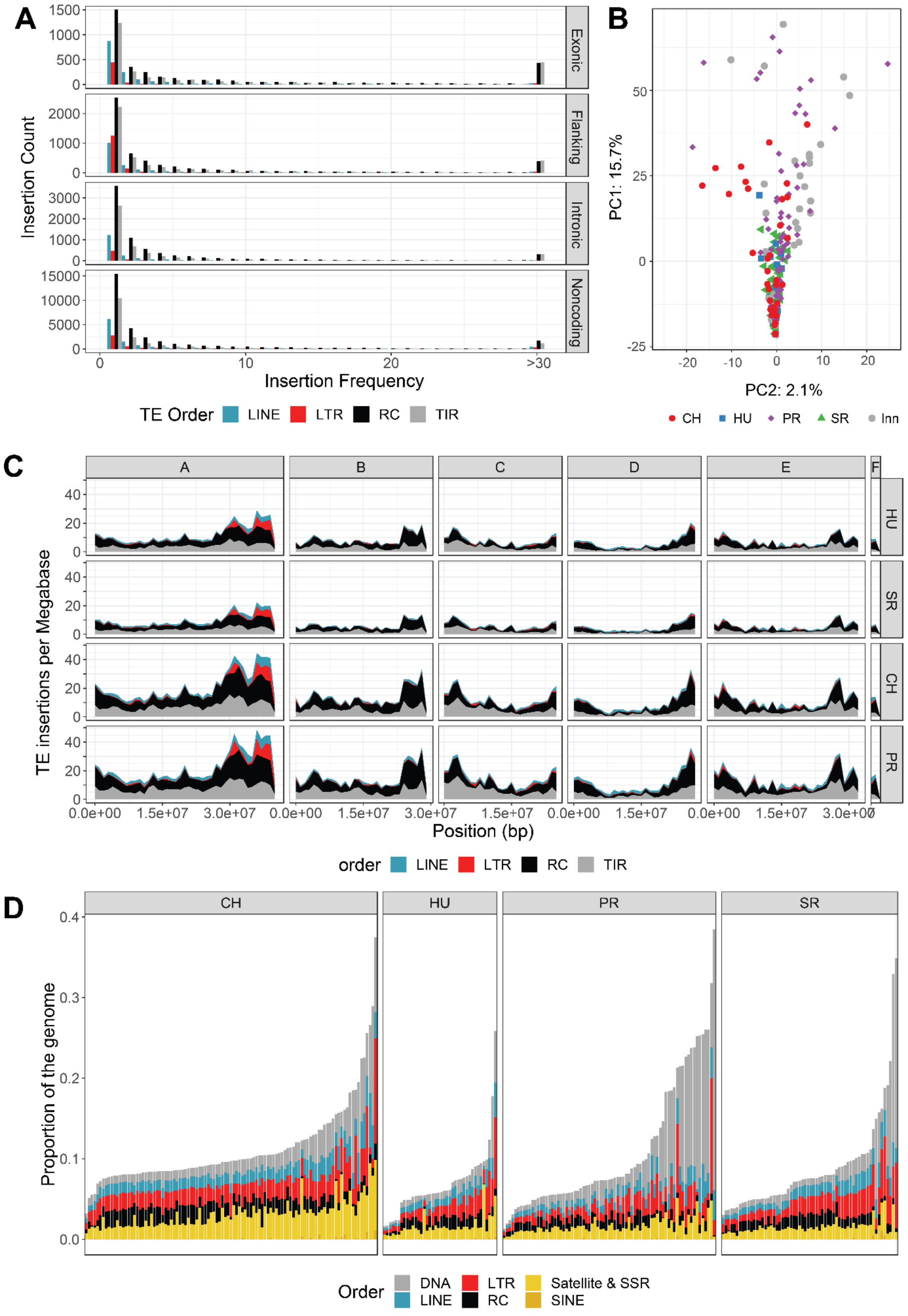

A
Insertion Count
Exonic
Flanking
Intronic
Noncoding
Insertion Frequency
TE Order LINE LTR RC TIR
B
PC1: 15.7%
PC2: 2.1%
CH HU PR SR Inn
C
A B C D E F
HU SR CH PR
TE insertions per Megabase
Position (bp)
order LINE LTR RC TIR
D
CH HU PR SR
Proportion of the genome
Order DNA LINE LTR RC Satellite & SSR SINE

**Supplementary Table 1:** Summary of *Drosophila innubila* fly's DNA collected and sequenced for this study, including summary of coverage for X chromosome, autosomes. Also contains SRA accessions for each strain. Coverage for different portions of Muller element B are shown when applicable.**Supplementary Table 2:** GLM for population genetic statistics in immune gene categories relative to the background for each population.

**Supplementary Table 3:** δaδi parameters estimated for each model, for each chromosome, for each population comparison. When parameters are not used, NA is shown. The significance from a likelihood-ratio test between the split population model (bottlegrowth_split) and the split population with migration model (bottlegrowth_split_mig) compared to the panmictic model (bottlegrowth) is shown when applicable. Abbreviations: nuB: Ratio of population size after instantanous change to ancient population size. nuF: Ratio of contempoary to ancient population size. m: Migration rate between the two populations (2*Na*m). T: Time in the past at which instantaneous change happened and growth began (in units of 2*Na generations). Ts: Time in the past at which the two populations split.

**Supplementary Table 4:** Summary of gene ontology enrichments for $F_{ST}$ in each population, separated by processes, components and functions.

**Supplementary Table 5:** Summary of GLM for elevated McDonald-Krietman statistics GO categories in *D. innubila*.

**Supplementary Table 6:** Summary of gene ontology enrichments for $D_{XY}$ in each population.

**Supplementary Data 1:** VCF file for SNPs in *D. innubila*, used in estimation of population genetic statistics and in GWAS.

**Supplementary Data 2:** Population genetic statistics calculated for each gene in *D. innubila* using VCFtools for each population.

**Supplementary Data 3:** McDonald-Kreitman statistics calculated for each gene in *D. innubila* using SnIPRE for each population.
